# A molecular switch for Cdc48 activity and localization during oxidative stress and aging

**DOI:** 10.1101/733709

**Authors:** Meytal Radzinski, Ohad Yogev, Yarden Yesharim, Esther S. Brielle, Ran Israeli, Rosi Fassler, Naomi Melamed-Book, Nadav Shai, Isaiah T. Arkin, Elah Pick, Tommer Ravid, Maya Schuldiner, Dana Reichmann

## Abstract

Control over a healthy proteome begins with the birth of the polypeptide chain and ends with coordinated protein degradation. One of the major players in eukaryotic protein degradation is the essential and highly conserved ATPase, Cdc48 (p97/VCP in mammals). Cdc48 mediates clearance of misfolded proteins from the nucleus, cytosol, ER, mitochondria, and more. Here we dissect the crosstalk between cellular oxidation and Cdc48 activity by identification of a redox-sensitive site, Cys115. By integrating proteomics, biochemistry, microscopy, and bioinformatics, we show that removal of Cys115’s redox-sensitive thiol group leads to accumulation of Cdc48 in the nucleus and consequently, results in severe defects in the oxidative stress response, mitochondrial fragmentation, and a decrease in ERAD and sterol biogenesis. We have thus identified a unique redox switch in Cdc48, which may provide a clearer picture of the importance of Cdc48’s localization in maintaining a “healthy” proteome during oxidative stress and chronological aging in yeast.

## Introduction

Protein quality control (PQC) is one of the most tightly regulated systems within the cell, from the initial steps of the polypeptide chain formation through proteasomal degradation^1^. Failure to address a misfolded protein (either by degradation, sequestration, or refolding pathways) not only leads to a loss of protein function, but may also trigger toxic protein aggregation within the cell^2^ and subsequent disorders^3^.

Cdc48 (p97/VCP in mammals) is a highly prevalent and conserved ATPase chaperone involved in regulating protein degradation processes^4, 5^. Cdc48 has diverse roles in various degradation pathways such as endoplasmic reticulum-associated degradation (ERAD)^6^, mitochondria-associated degradation (MAD)^7^, Rsp5-mediated degradation^8^, and more^9–11^. Cdc48 extracts and delivers misfolded proteins from their initial sites (e.g. the endoplasmic reticulum (ER) membrane following retrotranslocation) to the proteasome for degradation. Cdc48 has also been implicated in direct refolding activity, as bolstered by its two ATP-binding domains^12^ (Fig. 1A). Cdc48 is therefore a major regulator of proteasomal degradation pathways, necessary for delivery and recognition of misfolded client proteins. Accordingly, mutations throughout Cdc48 have also been found in several diseases (e.g. Paget’s Bone Disease, Amyotrophic Lateral Sclerosis (ALS))^13, 14^.

**Figure 1.**
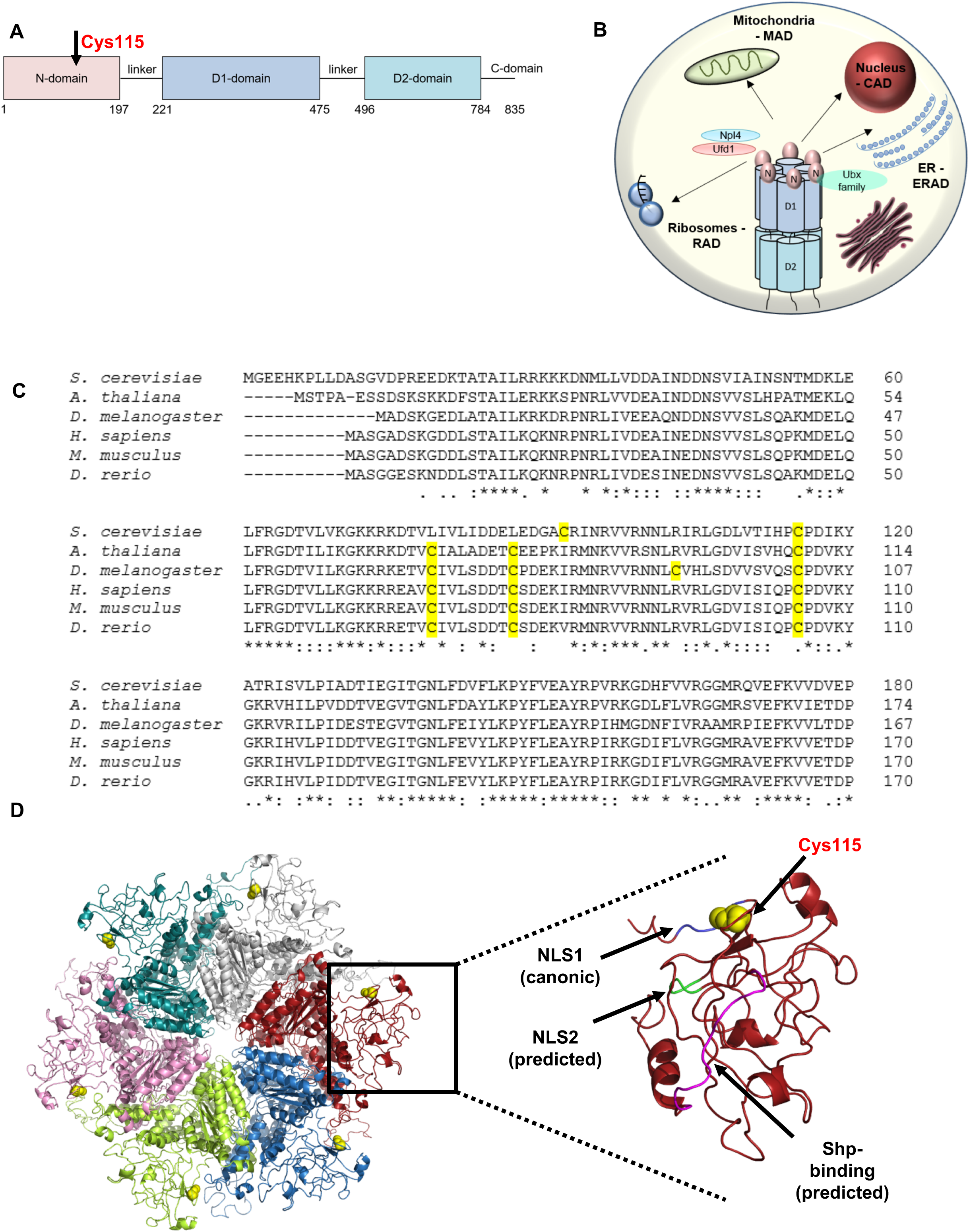
(A) Schematic representation of Cdc48’s sequence, with Cys115’s approximate location emphasized. (B) Scheme of Cdc48 localization and activity within the cell, including association with cofactors Npl4/Ufd1 and the Ubx family leading to degradation processes throughout the cell. (C) Multiple sequence alignment of Cdc48 homologs (yeast, Arabidopsis, Drosophila, human, mouse, and zebrafish, respectively), showing high alignment in the N-terminal domain with a single cysteine residue conserved between all organisms. All cysteine residues labeled in yellow. (D) Structural homology model of Cdc48 based on active p97 structure (PDB: 5FTN), zoomed to show N-domain. Cys115 is in yellow, the canonic NLS (NLS1) in blue, the predicted NLS (NLS2) in green, and the predicted Shp-binding region in magenta.

The N-terminal domain of Cdc48 serves as an anchor for multiple interactions with cofactors which define localization and activity of Cdc48^15, 16^. Two of Cdc48’s canonic cofactors are Npl4 and Ufd1, forming the Cdc48-Npl4-Ufd1 complex which is necessary for both substrate recognition and targeting of proteins for proteasomal degradation at the ER membrane (Fig. 1B)^17, 18^. At other organelles such as mitochondria, Cdc48 utilizes other cofactors (e.g. the Cdc48-Npl4-Vms1 complex mediates mitochondrial-associated degradation)^7^. Additional cofactors such as members of the Ubx-family are involved with Cdc48 in nuclear^19^ and membrane lipid formation^20^ pathways (Fig. 1B).

Cdc48’s N-terminal domain is relatively structurally flexible^21^, with such structural plasticity likely allowing multifunctional binding of diverse proteins. In addition, it was suggested that the motion of the N-terminal domain is coupled with its ATPase activity, and in turn is affected by mutations in the ATPase D1/D2-domains^14, 22–24^.

Previous research has identified yeast Cdc48 as a potential redox sensitive protein, which alters its oxidative status during chronological aging^25^. A redox proteomics analysis of yeast proteins revealed that a couple of days following the metabolic diauxic shift, almost 80% of thiol-containing proteins undergo sudden oxidation. Interestingly, a subset of proteins showed much earlier thiol oxidation during chronological aging, a day ahead of the general redox collapse that the rest of the proteins experience^25^. Cysteine 115 of Cdc48 is one such thiol. This follows an increasing number of recently identified “thiol switch” proteins, activity and structure of which depend on the oxidation of certain cysteine thiols. Post-translational modifications of specific thiol groups in this class of proteins allow a rapid response to environmental and physiological redox perturbations in cells^26, 27^. Identification of Cdc48 within the subset of early-oxidized proteins during aging raises the interesting hypothesis that Cys115 may have a regulatory role for Cdc48 function, thus poising Cdc48 as participating in preempting the redox collapse.

Here, we show that Cdc48 has a crucial role in maintaining redox homeostasis and that the N-terminal Cys115 is a unique thiol switch with implications for the global redox response, protein homeostasis, and cell survival. We found that mutation of Cys115 to Serine (Ser, which removes the redox-sensitive thiol) affects cell survival and growth, while also severely inhibiting cell growth during oxidative stress. Using confocal microscopy and proteomics, we identified strong localization of the mutated Cdc48 to the nucleus, providing new insights into Cdc48’s role during the oxidative stress response. We further identified impairment of ERAD and sterol biogenesis, alongside an increase in the unfolded protein response (UPR), a disruption of lipid homeostasis, and changes in mitochondrial morphology. Finally, we propose a model by which Cys115 contributes to Cdc48 localization and interaction with cofactors and membranes, thus defining Cdc48’s protein function during healthy conditions and oxidative stress.

## Results

### The Cys115 residue in Cdc48 regulates cell growth and survival during oxidative stress

Cdc48 is a highly conserved protein, with sequence identity maintained between a wide range of different homologs (Fig. 1C, Table S1). Despite the high identity between the different homologs, most of Cdc48’s cysteine residues are not well conserved between low and high complexity organisms. In the flexible N-terminal domain, only Cys115 remains conserved across a wide range of homologs (Fig. 1C). In addition to undergoing oxidation during the early stages of chronological aging^25^, Cys115 is also located on the surface-exposed region of the N-terminal domain based on high-homology modeling (using known p97 structures) (Fig. S1), with its exposure modeled to change according to N-terminal domain motion following ATP-binding^23^, alternating from a more buried pocket to an exposed segment following Cdc48 activation. Moreover, based on this modeling, Cys115 is physically located near both Cdc48’s canonical nuclear localization sequence (NLS) and a predicted NLS (Fig. 1D)^28^, as well as a region modeled to be a Ubx-motif-binding site^29^ (Fig. 1D). Together, these place Cys115 in a clear position to be a major redox regulator, with the potential to influence Cdc48’s protein-protein interactions across a wide range of functions.

To investigate the effect that the oxidation of Cys115’s thiol may have on redox and protein homeostasis at large, we introduced a point mutation, replacing the active thiol with an oxidation-insensitive OH group through mutation of the cysteine to a serine residue (C115S) (Fig. 2A). Both FLAG-tagged wild type (wt) and Cdc48-C115S were expressed on plasmids on the background of a deletion in the essential, endogenous CDC48 gene.

**Figure 2.**
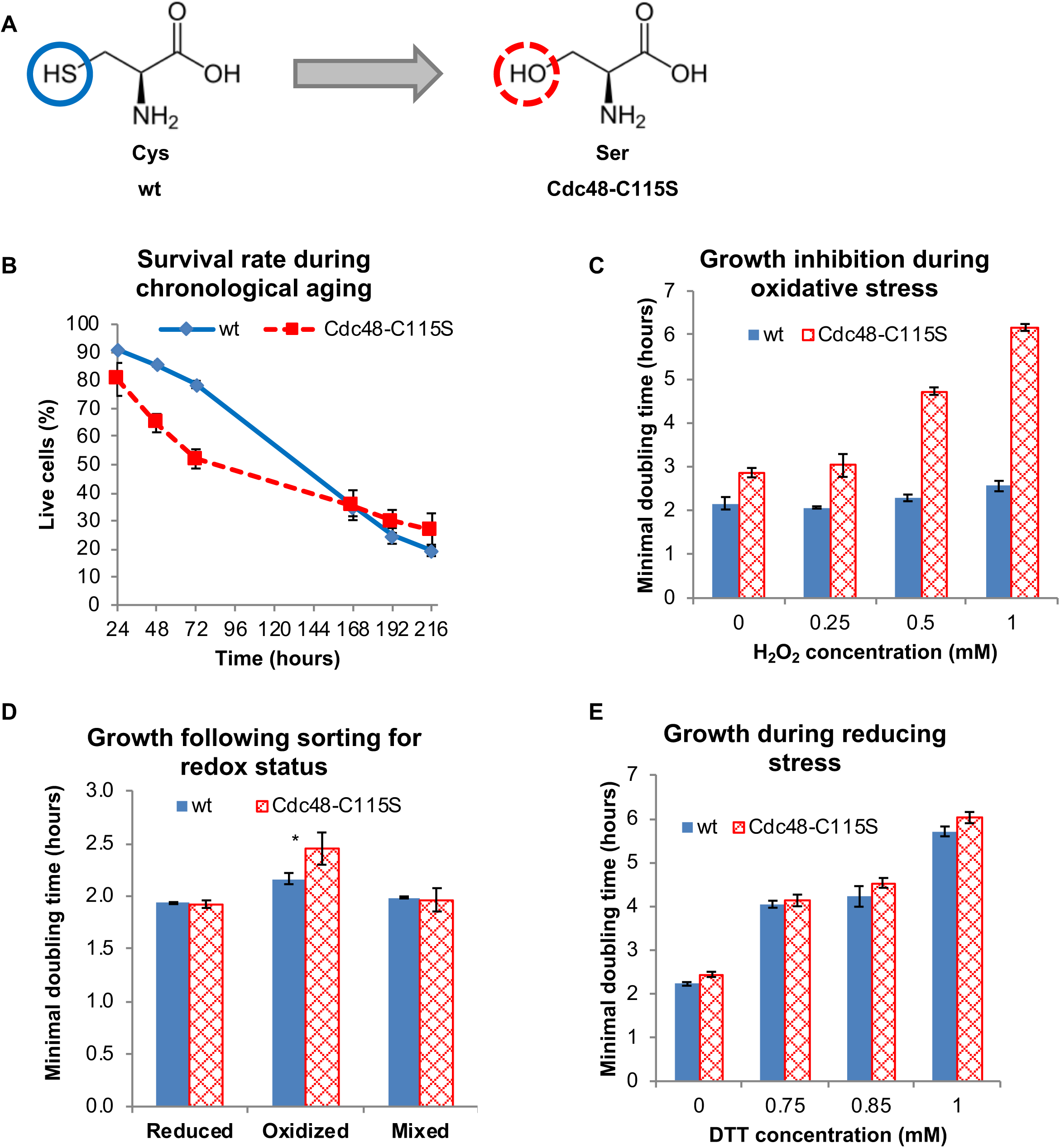
Cdc48-C115S impairs growth. (A) Schematic representation of Cdc48-C115S mutation. (B) Survival of living cells expressing either the wt or Cdc48-C115S variant. The Cdc48-C115S strain has decreased vitality as compared to the wt during chronological aging. (C-D) Cdc48-C115S has severe growth inhibition (higher doubling times) during oxidative stress of varying degrees (C) or after isolation of natively oxidized and reduced subpopulation of cells (D). P-values for treatments all less than 0.05. The sorting was based on an in-vivo expressed roGFP sensor, which changes its ratiometric fluorescence with oxidation^70^. Oxidized, reduced, and unsorted (mixed) cultures were diluted and doubling time was calculated for each subpopulation. (E) Reducing stress (using varying concentration of DTT) does not affect the Cdc48-C115S growth.

To validate the oxidation profile of Cys115 in the newly designed strains, we decided to investigate if indeed this cysteine undergoes oxidation in vivo as previously shown by the Jakob lab^25^. We conducted a differential thiol-trapping analysis of Cdc48 using two different alkylation reagents (iodoacetamide (IAM) and N-Ethylmaleimide (NEM)), followed by mass spectrometry (Fig. S2). Cdc48 was enriched via co-immunoprecipitation using a FLAG-tag from cells grown for 24h and 48h, in three biological replicates. After denaturation, reduced cysteines were irreversibly labeled by IAM (Fig. S2A). We then reduced all oxidized cysteines by TCEP and alkylated all newly reduced cysteines by NEM (Fig. S2A). After trypsin digestion and peptide analysis, we quantified differences in intensities between peptides containing either NEM- or IAM-modified Cys115, which represent differences between reduced and reversibly oxidized Cys115 in-vivo. In line with the previous redoxome analysis, we also found that Cdc48-Cys115 underwent oxidation with age; while the 24h-aged culture harbored both reduced and oxidized Cys115 with a preference for reduced, the 48h culture mainly contained reversibly oxidized Cys115 (Fig S2B-D). It is important to note, that this analysis cannot identify irreversibly oxidized modification, which might exist with increased levels of oxidants in cells.

Next, we assessed the overall vitality of cells expressing either the wt or Cdc48-C115S variant during chronological aging. We found that, when grown on synthetic medium supplemented with casein, the mutant strain had a decreased survival rate as compared to the wt from the early stationary phase (24h) through to late stages of chronological aging (Fig. 2B). Interestingly, the largest difference in vitality was observed at day 2 (48h) and day 3 (72h), which are associated with increased cellular oxidation, as well as the Cys115 oxidation itself. Growth was also impaired to a significant degree on standard minimal medium (Fig. S3A). We also noticed a slightly slower minimal doubling time (growth rate) during the logarithmic phase, though this result was not always statistically significant, most probably due to small variabilities in growth conditions or initial cell number (Fig. S3B).

Having identified changes in growth and cell survival in Cdc48-C115S during normal conditions, we next examined the effect of this point mutation on the cellular stress response, including oxidation and reduction stresses. We conducted a series of growth tests to assess the response of cells expressing Cdc48-C115S to oxidative stress and found a significant decrease in growth as compared to the wt (Fig. 2C-D). Whereas varying concentrations of hydrogen peroxide were largely tolerated by the wt strain, Cdc48-C115S’s growth was strongly inhibited (Fig. 2C), resulting in an elevated minimal doubling time.

To further examine such a strong effect of the Cdc48-C115S mutation on the sensitivity of cells to oxidative stress, we decided to examine potential differences in the growth of cells naturally different in their redox status, under physiological conditions. We applied flow cytometry to isolate cells with distinct redox statuses based on the differential fluorescence ratios at 405 and 488nm of the Grx1-roGFP2 sensor^30^ expressed in either wt or Cdc48-C115S cells. While cells with reduced cytosolic levels had a similar growth profile, oxidized cells expressing the Cdc48-C115S variant had a longer division rate (Fig. 2D). This is in line with our previous observation regarding the significant decrease in longevity during chronological aging (Fig 2B), which is associated with the natural accumulation of oxidants.

Interestingly, converse treatment with the reductive agent di-thio threitol (DTT) led to no significant difference between the strains (Fig. 2E). This suggests an oxidation-specific alteration to Cdc48 functionality following mutation of Cys115, rather than a general stress response. Additional experiments comparing growth of Cdc48-C115S during heat stress (37°C) and on low (0.5%) glucose medium revealed a similar lack of phenotype (Fig. S3C, S3D), strengthening our hypothesis that cells expressing Cdc48-C115S are uniquely sensitive to oxidation as a result of a single missing thiol group in the Cdc48 protein.

### Cdc48-C115S leads to abnormal mitochondrial morphology upon oxidative stress

The increased sensitivity of Cdc48-C115S cells to oxidative stress led us to hypothesize that redox sensitivity might be rooted in mitochondrial activity. To test mitochondrial morphology, we used a green fluorescent protein (GFP) variant targeted to the mitochondrial matrix^31^. While both the wt and Cdc48-C115S strains had fully intact mitochondria under normal growth conditions during log phase, we observed a significant difference during oxidative stress (Fig. 3). Under treatment with 1mM H_2_O_2_, we noted only minor perturbations to mitochondrial morphology in the wt, with most cells maintaining intact mitochondria as in the untreated samples. However, an overwhelming majority of the Cdc48-C115S cells displayed damaged, probably fragmented, mitochondria. Following results in the literature by which Cdc48 is known to localize to the mitochondrial membrane during oxidative stress^7^ as well as our earlier results, this strongly suggests that the Cdc48-C115S is unavailable for recruitment to the mitochondria during oxidative stress and as such leaves mitochondria more sensitive to oxidative damage.

**Figure 3.**
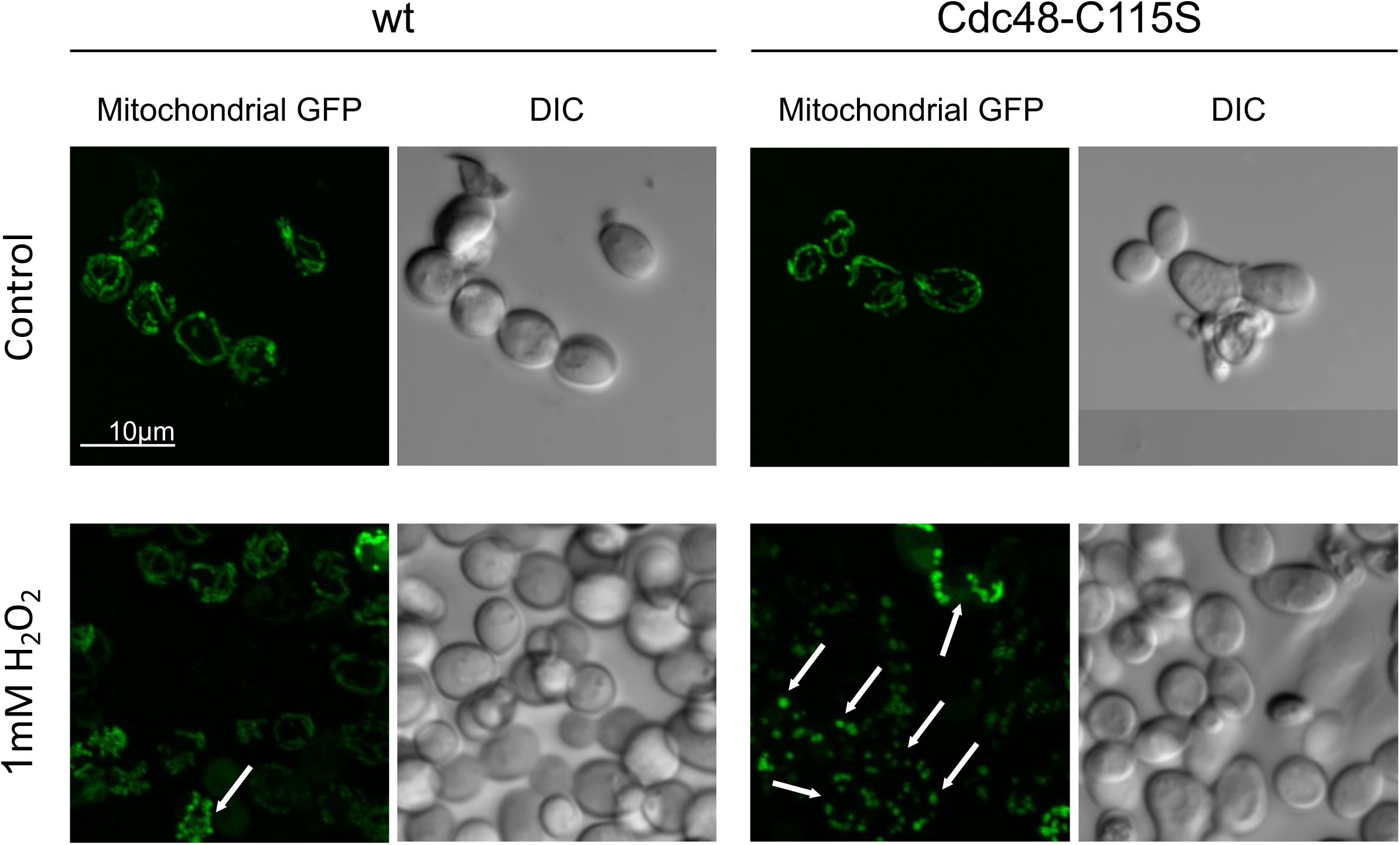
Confocal microscopy Z project of fluorescence stacks in the wt and Cdc48-C115S cells expressing mitochondrial targeted GFP during the logarithmic phase, under normal and oxidative conditions. Punctate and fragmented mitochondria are indicated by white arrows.

### Cys115 has a mild impact on redox homeostasis in yeast

Intrigued by peroxide sensitivity and changes in mitochondria shape upon oxidative stress, we conducted a global proteomic analysis of the wt and Cdc48-C115S expressing cells during chronological aging (late log phase (24h) and post-diauxic shift phase (48h)). This proteomic analysis identified 1731 proteins at 24h and 1716 at 48h (out of either 3 or 4 repeats, Tables S2, S3) and pointed to relatively few differences in protein profile between the strains (Fig. S4, Tables S4, S5). While there were few upregulated proteins in Cdc48-C115S cells (with no single pathway identified as having significant enrichment), we noted a relative decrease in general oxidoreductase activity and oxidative stress-associated proteins at both 24h and 48h as compared to the wt (Fig. S4C-F). Notable among these was the protein Whi2, a general stress responsive protein that has been found to play a role in mitophagy^32^ and was not expressed in the Cdc48-C115S strain at either time point (Tables S4, S5). Moreover, known redox homeostasis proteins protecting cells against oxidative stress (particularly as induced by hydrogen peroxide) were significantly downregulated at 24h upon mutation of Cys115 to Serine, including the catalase Ctt1, Trx2, and Sod1^33–35^ (S4, Tables S4, S5).

Taken together, these points towards a slight downregulation of proteins involved in the oxidative stress response in the Cdc48-C115S, which can explain sensitivity to oxidative stress and decreased growth during chronological aging.

### The Cdc48-C115S mutation preferentially localizes Cdc48 to the nucleus

To better understand the observed phenotypic changes between the wt and the Cdc48-C115S strains, we expressed the two Cdc48 variants fused with a GFP. Since Cdc48 is an essential gene, we performed a plasmid-shuffle between these Cdc48-GFP plasmids into our *Δcdc48* strains with either the wt or mutant Cdc48-FLAG plasmid (Fig. 4A). Interestingly, it was relatively easy to obtain strains with the Cdc48-C115S-GFP fusion only, while cells transformed with the wt form of the plasmid kept either both plasmids (fused with FLAG and GFP, respectively) or only the FLAG tagged protein. Therefore, we compared GFP expression in wt and Cdc48-C115S strains with both GFP- and FLAG-tagged Cdc48 that may include chimeric Cdc48 or altered expression (henceforth referred to as GFP* strains), alongside GFP expression in the Cdc48-C115S-GFP-only strain (referred to as GFP).

**Figure 4.**
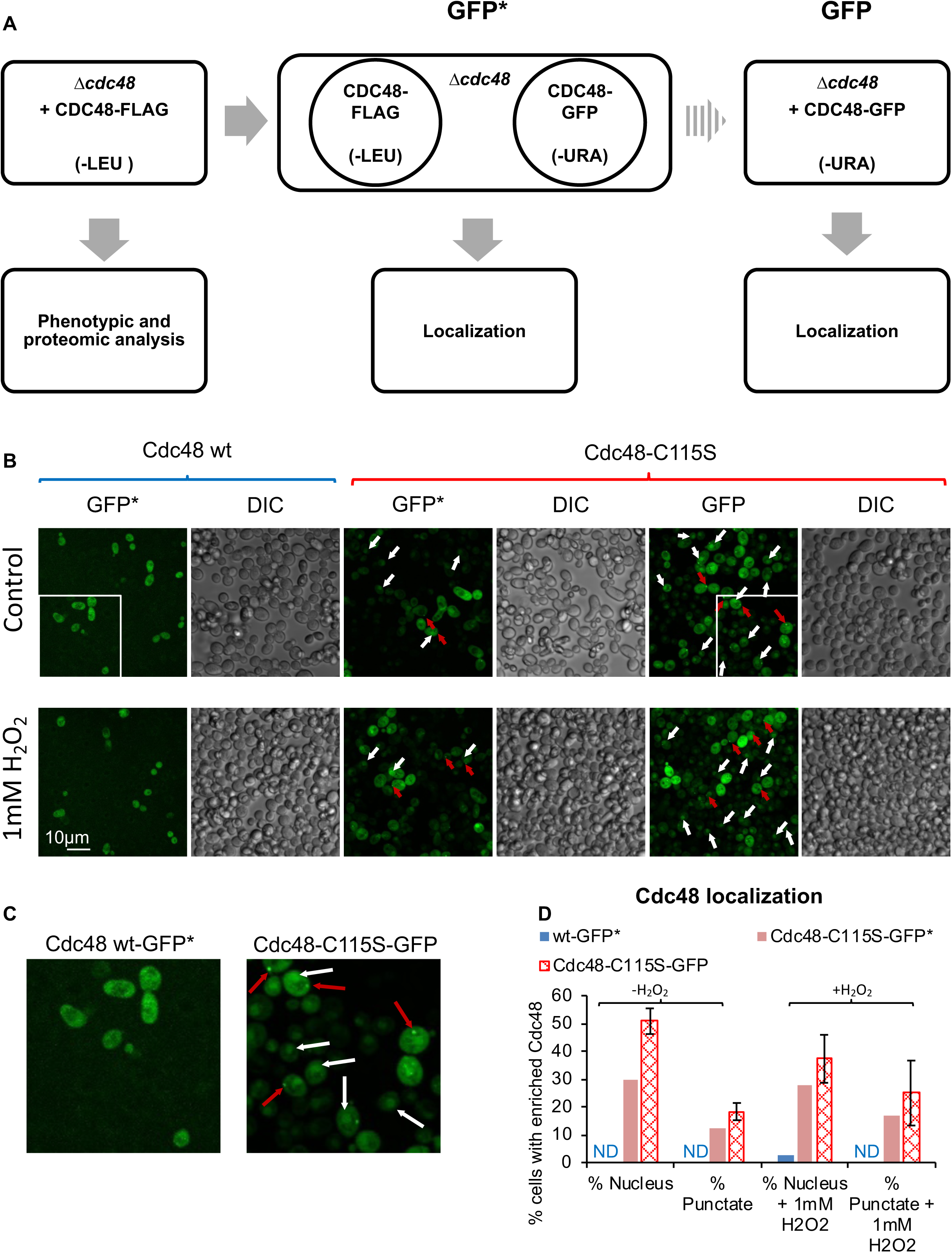
(A) Schematic representation of strains used. *Δcdc48*::CDC48-FLAG strains of either Cdc48 variant (wt or Cdc48-C115S; -LEU selection) were prepared and used for phenotypic and proteomic analysis. These strains were also transformed with the corresponding Cdc48-GFP plasmid (-URA selection). Strains which maintained growth on both selective mediums were labeled GFP* (i.e. expression of both Cdc48-FLAG and Cdc48-GFP), while those with selection on -URA alone were labeled GFP (i.e. expression of the Cdc48-GFP alone). Both GFP* and GFP strains were then taken for imaging to assess localization. (B) Expression of GFP-tagged Cdc48 using confocal microscopy. Cells expressing Cdc48 through both -FLAG and -GFP forms are labeled GFP*. GFP expression alone is labeled as such. White arrows indicate nuclear localization while red arrows indicate punctate localization. (C) Zoomed-in fields as labeled in (B). (D) Quantification of Cdc48 localization per confocal microscopy images as shown in (A) and S5. ND – not detected. Two fields with a total of 49 and 43 GFP-positive cells analyzed for wt-GFP* untreated and with 1mM H_2_O_2_ (respectively). Two fields with a total of 127 and 147 GFP-positive cells analyzed for Cdc48-C115S-GFP* untreated and with 1mM H_2_O_2_ (respectively). Three fields with a total of 363 and 366 GFP-positive cells from two experiments analyzed for Cdc48-C115S-GFP untreated and with 1mM H_2_O_2_ (respectively).

Using confocal microscopy, we noted a clear shift in Cdc48-C115S localization to the nucleus (Fig. 4B-C, S5, white arrows), as well as increased localization to puncta in various parts of the cell (Fig. 4B, 4C, S5, red arrows). Both variants of Cdc48-C115S (GFP* and GFP) had elevated levels of nuclear localization (Fig. 4B-D, S5), though it is notable that the stronger shift exists in Cdc48-C115S-GFP, which consists of one copy of the Cdc48 gene, similar to the Cdc48-C115S-FLAG construct. Expression of the Cdc48-C115S protein neither alone (GFP) nor alongside Cdc48-C115S-FLAG (GFP*) changed drastically during oxidative stress, with localization remaining nuclear with some puncta. However, it is interesting that the ratio between cells with puncta-localized Cdc48 and cells with nuclear-localized seems to decrease somewhat upon oxidative stress (Fig. 5D). Furthermore, Cdc48-C115S-GFP cells had a wider range of puncta-localized Cdc48 (as seen in the high standard deviation) during oxidative stress, which may point to broader localization heterogeneity during stress conditions.

**Figure 5.**
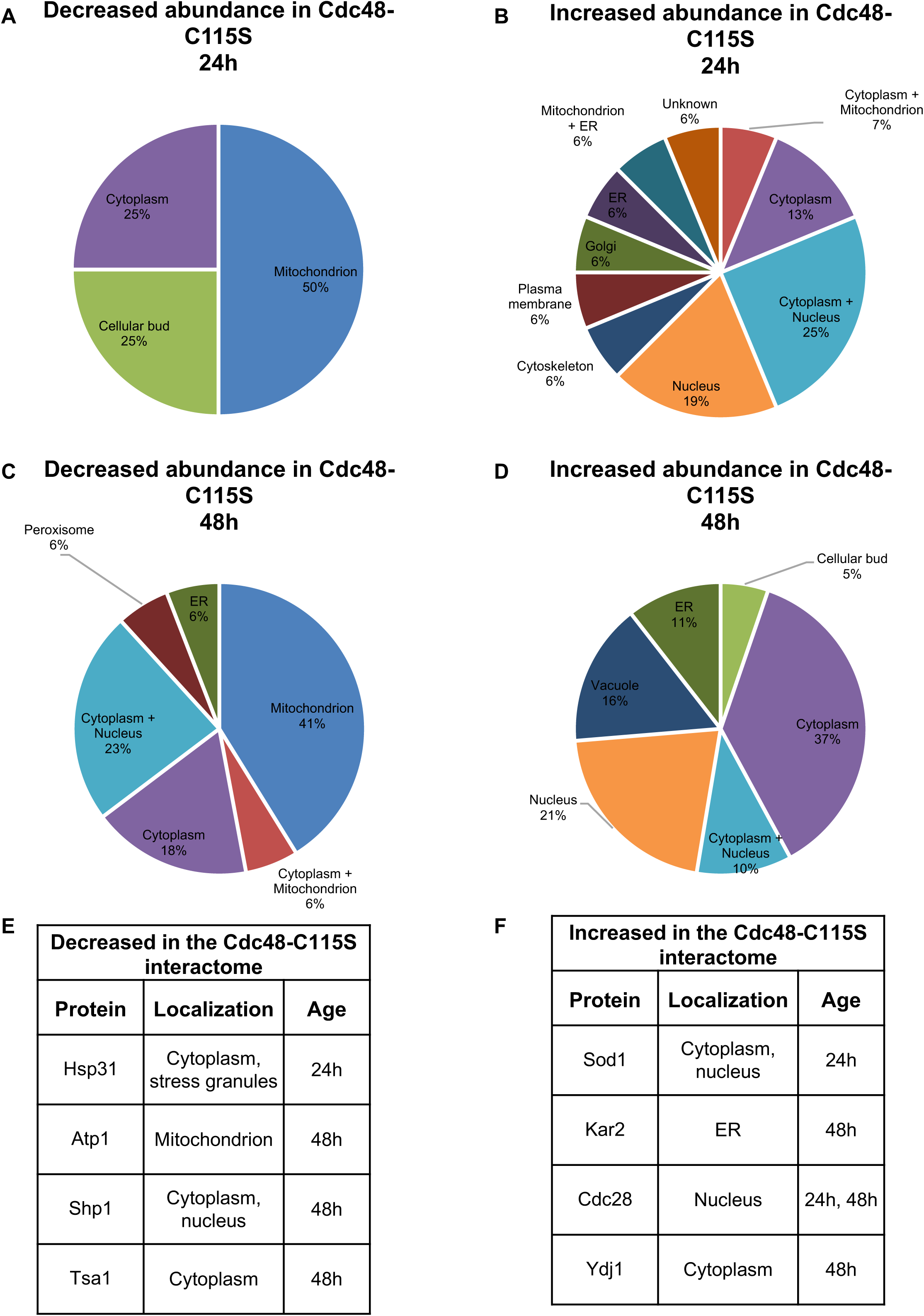
Changes in interactome of Cdc48 upon the Cdc48-C115S mutation. (A-B) Localization of proteins from Table S8 with relatively decreased (A) or increased (B) abundance in Cdc48-C115S at 24h. (C-D) Localization of proteins from Table S9 with relatively decreased (C) or increased (D) abundance in Cdc48-C115S at 48h. Cdc48-C115S is colocalized with markedly more nuclear proteins and fewer mitochondrial proteins. (E-F) Select proteins of interest from Tables S8, S9.

It is important to note that wt Cdc48-GFP expression was consistently low across all experiments and samples, requiring significant changes to the microscope fluorescent readout. Therefore, we cannot compare GFP intensity between the different samples, nor can we accurately gauge wt Cdc48 localization within those cells due to the weak expression. However, we may compare our results to those detailed in the literature, noting that while Cdc48 is known also to localize to the nucleus (in addition to the cytosol and mitochondria^7, 36, 37^), high levels of nuclear localization are typically found under specifically induced conditions (e.g. substrate solubility)^38, 39^. Thus, we found that the Cdc48-C115S mutation leads to preferential Cdc48 localization to the nucleus at significantly higher rates than the wt without applying any induced conditions.

It is tempting to speculate that Cdc48-C115S serves as a model for a constitutively nuclear form of Cdc48, which can be used to investigate role of the Cdc48 localization within the cell and its impact on cellular survival during stress conditions.

### Mutation in Cys115 alters Cdc48 interactome, reflecting differences in localization and interaction with antioxidants

Intrigued by the observed changes in Cdc48 localization following mutation of Cys115 alongside minor changes in the global protein profile, we decided to examine interactome of the wt and Cdc48-C115S variants in cells at 24h and 48h growth (before and after the diauxic shift) through co-immunoprecipitation using FLAG-tagged Cdc48. As a control, we conducted an additional pull-down of the background wt strain (BY4741) without the Cdc48-FLAG plasmid in order to characterize nonspecific protein binding. The rationale behind this extensive analysis was to investigate potential changes in the Cdc48’s interacting proteins which can reflect cellular localization of the Cdc48 variants and explain the differences in stress response as well the mitochondrial morphology. This analysis identified 885 and 858 proteins total at 24h and 48h (respectively) (Tables S6, S7). To identify Cdc48’s specific binding partners at 24h and 48h, we removed all non-specific binders identified in the non-FLAG wt strain. Comparing the normalized protein abundances (LFQ) of the remaining specifically interacting proteins (between wt or Cdc48-C115S), we identified striking differences in the interactomes of the two strains, with a total of 22 and 37 significantly differentially binding proteins at 24h and 48h, respectively (Tables S8, S9).

In line with the imaging experiments, the wt and the Cdc48-C115S proteins bind proteins located in different organelles (Fig. 5). Specifically, we identified a strong decrease in the number of mitochondria-associated proteins in the Cdc48-C115S pull-down (Fig. 5A, C), alongside a stark increase of nuclear proteins (Fig. 5B, D). These support our previous observation of a change in Cdc48 localization following Cdc48-C115S mutation and sensitivity of mitochondria to oxidative stress. Moreover, the interactome profile of Cdc48-C115S had a higher abundance of cytoskeleton and membrane-associated proteins, as well as of vacuolar proteins (Table S8, S9).

Upon examining the subset of significantly changed proteins between the wt and Cdc48-C115S interactome, we found that significant portions are involved in pathways associated with canonic Cdc48 functions: ERAD and the stress response (Fig. 5E, F, Tables S8, S9). We identified a decrease in abundance of the stress response chaperone Hsp31 (the yeast homolog of the Parkinson’s Disease associated human DJ-1^40, 41^) in the Cdc48-C115S strain at 24h, as well as oxidative stress associated proteins such as the thioredoxin Trx1 and the peroxiredoxin Tsa1 at 48h, strengthening our understanding that Cdc48-C115S may have altered redox regulation. Importantly, we also identified a decreased association between the mutant Cdc48 and the Ubx-family member Shp1/Ubx1 at 48h. Shp1 is a known cofactor of Cdc48 associated with ERAD, nuclear membrane functions^42^, and autophagy^13^. Shp1 has been suggested as an antagonist of Ufd1, interacting with Cdc48 independently of the Ufd1-Npl4 complex and simultaneously with ubiquitylated substrates^5^. Its decreased abundance in the Cdc48-C115S interactome is a strong indicator of a shift in canonic Cdc48 activity, though the precise mechanism is still unclear. Meanwhile, the abundance of additional proteins associated with ERAD rather than with direct Cdc48 interactions (e.g. ubiquitin conjugating enzymes, Kar2/BIP, Ydj1/Hsp40) was relatively decreased in the Cdc48-C115S interactome profile, pointing towards altered protein quality control regulation in the ER or association with ERAD.

Following this analysis, we assumed that different colocalizations could alter the relative influence of the background interacting proteins (i.e. FLAG bead-binding proteins) removed during the strict analysis of the wt- and Cdc48-C115S-binding proteins as previously shown. Therefore, we directly compared all identified proteins in the wt and Cdc48-C115S pull-downs, without removal of the FLAG bead-binding proteins as is commonly done in comparison of multiple pull-down experiments, varying in conditions and treatments^43, 44^. The rationale behind this analysis is to compare direct and indirect interactors of Cdc48, which might provide a broader picture of the Cdc48 localization. By this analysis, we identified 141 and 155 proteins with significantly different abundance between the wt and the Cdc48-C115S interactome at 24h and 48h, respectively (Tables S10, S11). It is important to note, that the abundance of significantly different binders of Cdc48-C115S and wt identified in the pull-down experiment did not show altered abundance in the global proteomic analysis. Thus, differences in the interactome profile of the Cdc48 variants are due to differences in binding with either wt or mutant Cdc48 variants rather than due to their expression levels.

Similarly to the more strict analysis described previously, we identified a relative decrease in expression of antioxidants, redox-associated proteins, and chaperones at 48h, such as catalase (Ctt1), ATP-independent chaperone (Hsp26), and Hsp31 in the Cdc48-C115S strain at both 24h and 48h (Fig. S6C, Fig. S6E), as well as an additional decrease in antioxidants such as thioredoxins (Trx1/Trx2), glutaredoxins (Grx2), and peroxiredoxins (Tsa1) (Tables S10, S11). Most probably, these are indirect interactions with Cdc48, which suggest perturbation among Cdc48 downstream substrates. This decrease in Cdc48 redox homeostasis binders correlates strongly with the oxidative stress sensitivity of the Cdc48-C115S strain, as previously described (Fig. 2D).

Generally, we found striking differences between the two strains (Fig. S6A, S6B), noting a strong shift in abundance of RNA-binding proteins, translation-associated and ribosomal proteins in Cdc48-C115S at 24h, including the ribosomal quality control-associated Rqc2 (a known cofactor of Cdc48 in ribosome-associated degradation^45^) and ribosome biogenesis proteins such as Rpf2 and Enp2 (Fig. S6D). At 48h, the differences were even more pronounced, with an increased expression of 13 proteins belonging to the 40S or 60S ribosomal subunits (Fig. S6F). This accumulation of ribosomes, which most probably are not direct interactors of Cdc48 itself, might indicate potential proteotoxic stress conditions, including the accumulation of misfolded proteins or oxidative stress^30^.

We may thus speculate that Cdc48-C115S’s sensitivity to oxidative conditions extends beyond exogenously induced stress, due to mislocalization of Cdc48-C115S in cells which leads to a decreased number of Cdc48 molecules in the cytosol and consequently a decreased ability of cells to degrade oxidatively damaged mitochondrial proteins. Thus, mitochondria in the Cdc48-C115S mutant are sensitive to mild perturbation of the redox status, resulting in mitochondrial alteration and likely dysfunction.

### Impact of Cysteine 115 on the dynamics of Cdc48’s N-terminus and NLS domain

Next, we sought to understand the structural basis of the tremendous impact of Cysteine 115 toward Cdc48’s localization and cellular oxidation. Due to the difficulty in assessing Cdc48’s structure generally and the N-domain more specifically, we conducted molecular dynamics (MD) simulations to monitor potential changes in the Cdc48’s structure upon either removal or oxidation of the C115 thiol group. To do so, we used the previously described model of the Cdc48 N-terminal domain (1-221aa) (Fig. 1, S1) as the wt structure. In addition to the wt structure, we created three *in silico* mutants: Cdc48-C115S, as well as oxidized forms of Cdc48-C115 to produce either sulfenic acid (C115-SO) or sulfinic acid (C115-SO_2_). We performed 100ns production MD simulations of each of these four models and compared the fluctuations of each residue.

In order to determine Cdc48 stability for each of the models, we used FoldX to calculate the folding energy for each frame between 10 and 100 ns of the production simulation. Cysteine oxidation has little effect on the global energy of Cdc48, whereas the serine mutation, destabilizes the protein (Fig. S7A).

Analysis of structural fluctuations during the MD progression showed that the Cdc48-C115S mutation leads to increased fluctuations of the NLS domains (RRKKKDN, 28-34aa and KKRKDT –71-76aa) (Fig. S7B-C, blue). In contrast, oxidation of Cys115 either has no impact (in NLS-2) or even decreases structural fluctuations (in NLS-1), leading to a more compact conformation and NLS protection (Fig. S7B-C, red).

Based on molecular dynamics and redox proteomic analysis (Fig. S2), we may then hypothesize that during chronological aging associated with an increase in oxidants, there is accumulation of Cdc48 with oxidized Cys115 residues. Oxidized Cys115 stabilizes the Cdc48 N-terminus and prevents, at least partially, its migration to the nucleus. Thus, accumulation of Cdc48 in the cytosol provides an anti-oxidative defense for cells by mediating degradation of damaged proteins in the cytosol, mitochondria, and ER.

### Cdc48 association with ERAD is mediated by Cys115

To more fully understand the differences in our phenotypic, proteomic and interactomic analyses induced by the Cdc48-C115S mutation, we focused on Cdc48’s involvement in canonic, ER-associated pathways (Fig. 6A). We first characterized Cys115’s role in ER-associated degradation (ERAD)^46^, measuring induction of the unfolded protein response (UPR) (induced when ERAD is not optimally functioning) using a reporter in which GFP is expressed under a UPR element (UPRE)^48^. We found that the Cdc48-C115S strain exhibited a higher normalized expression of the UPRE-GFP as compared to the wt during stationary phase, suggesting that Cys115 is involved in mediating Cdc48’s role in ERAD, most probably by mislocalization to nucleus (Fig. 6B). We further examined the effect of Cdc48-C115S on the degradation kinetics of one an ERAD substrate, the ER luminal protein KHN^49^, which requires functional Cdc48 for its retrotranslocation and degradation^50^. Mutation of Cdc48-C115S slightly slows KHN degradation over time as compared with the wt (Fig. 6C-D, representative data; Fig. S8), again pointing towards an impairment in Cdc48’s role in ERAD. These results further correspond with a decrease in abundance of the ERAD associated Cdc48’s cofactor Ubx1 (Shp1), as well as increased levels of Kar2 and Ydj1 in the Cdc48-C115S interactome set (Tables S8, S9). The UPR element is found in the promoters of both Kar2 and Ydj1, thus aligning with the increased UPR measured by our reporter.

**Figure 6.**
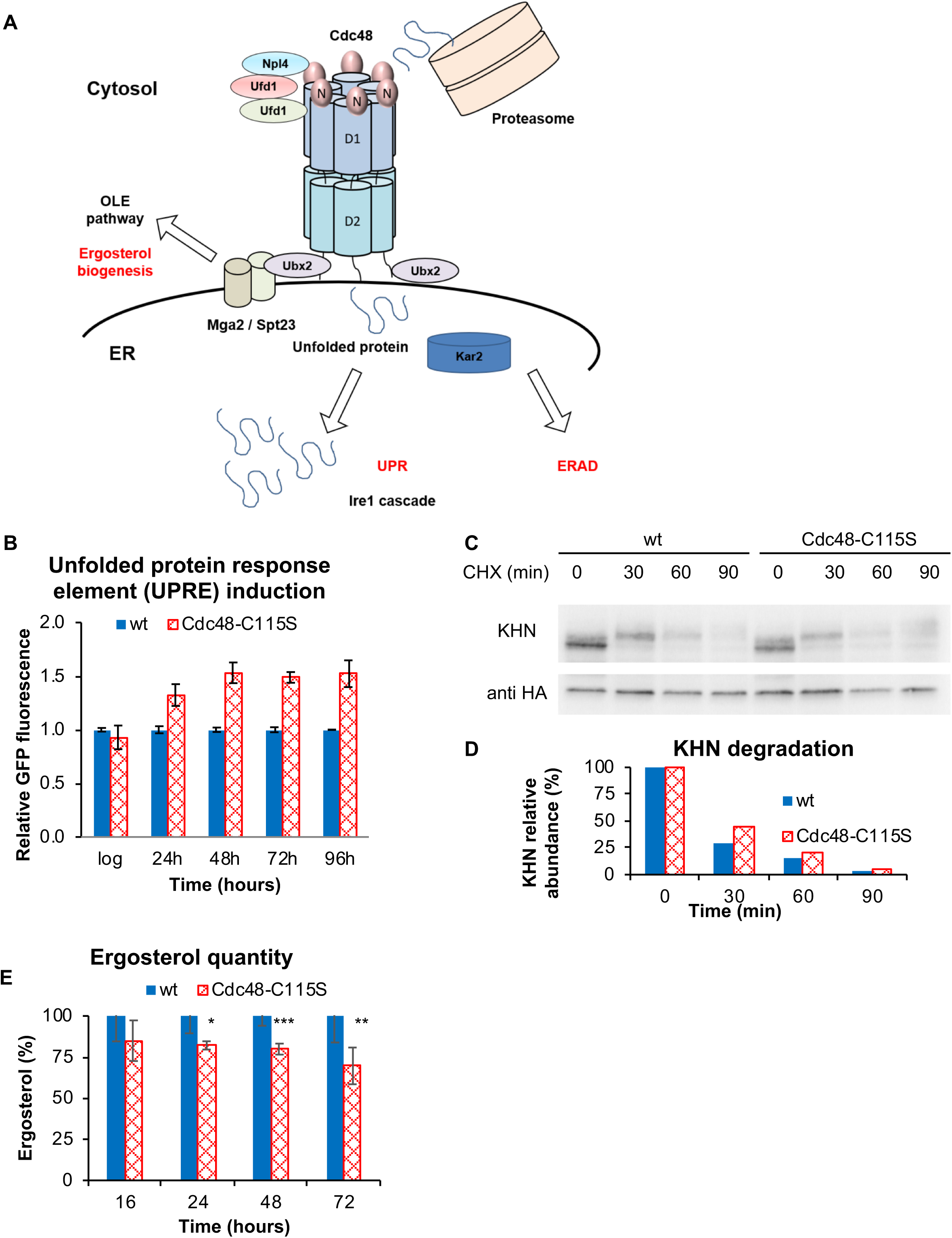
Cdc48 interaction with ER. (A) Schematic representation of Cdc48 interaction with different ER-associated pathways, including activation of the OLE and UPR pathways as mediated by proteins such as Ubx2. (B) Expression of the GFP-tagged unfolded protein response element, normalized to wt expression during chronological aging. Cdc48-C115S has a significantly higher fluorescence relative to the wt, i.e. an elevated UPR. (C-D) Representative pulse-chase experiment demonstrating Cdc48-C115S’s impaired degradation of the HA-tagged ERAD substrate, KHN. In (C), anti-HA blot of the HA-tagged degradation substrate KHN, while (D) quantifies the values from (C). (E) Ergosterol quantity normalized to wt expression during chronological aging. Cdc48-C115S has an increasingly reduced quantity of ergosterol compared to the wt.

Furthermore, Cdc48 is linked to its substrates through adaptor proteins, including a family of proteins with a Cdc48-binding UBX domain. Among other Ubx proteins, the Cdc48-Ubx2 interaction has been identified as a key player in the regulation of membrane lipid biosynthesis, required for cleavage of the 120 kDa precursor of Spt23 and Mga2^20, 51, 52^. These result in the formation of p90 transcription activators of the target gene *OLE1*, encoding the fatty acid desaturase, as well as additional *ERG* genes, encoding the early components of the ergosterol biosynthesis pathway (mevalonate pathway)^53–55^. Using the spectrometric method, we first quantified ergosterol levels in various control strains (including BY4741 wt α and a strains, Ubx-family deletion strains, and a mutation in the key ubiquitin E3 ligase *rsp5-1*) (Fig. S9A), finding that the mutants – *Δubx2* and *rsp5-1* more significantly – had decreased ergosterol levels, as expected. We then quantified ergosterol in the wt and Cdc48-C115S strains, finding that Cdc48-C115S had decreased ergosterol during chronological aging (Fig. 6E). We further compared this to a deletion of *ERG6* during oxidative stress, finding its ergosterol level comparable to that of the Cdc48-C115S (Fig. S9B). Cdc48-C115S’s reduced ergosterol levels suggest reduced functionality of the mutant Cdc48 involvement in ergosterol biogenesis, possibly through altered interaction with Ubx2.

To summarize, our study suggests that localization of Cdc48 is mediated by a highly conserved Cysteine 115 and affects its activity in protein quality control during oxidative stress (Fig. 7). We propose that this role is rooted in the redox status of Cysteine 115, which undergoes reversible modification upon exposure to an oxidative environment^25^ (Fig. S2). Use of the Cdc48-C115S mutant opens a door to study a role of the mislocalization of Cdc48 in the cell, and the importance of its presence in the cytoplasm during oxidative stress conditions, in order to sustain the health of cellular proteins as well as organelles (Fig. 7).

**Figure 7.**
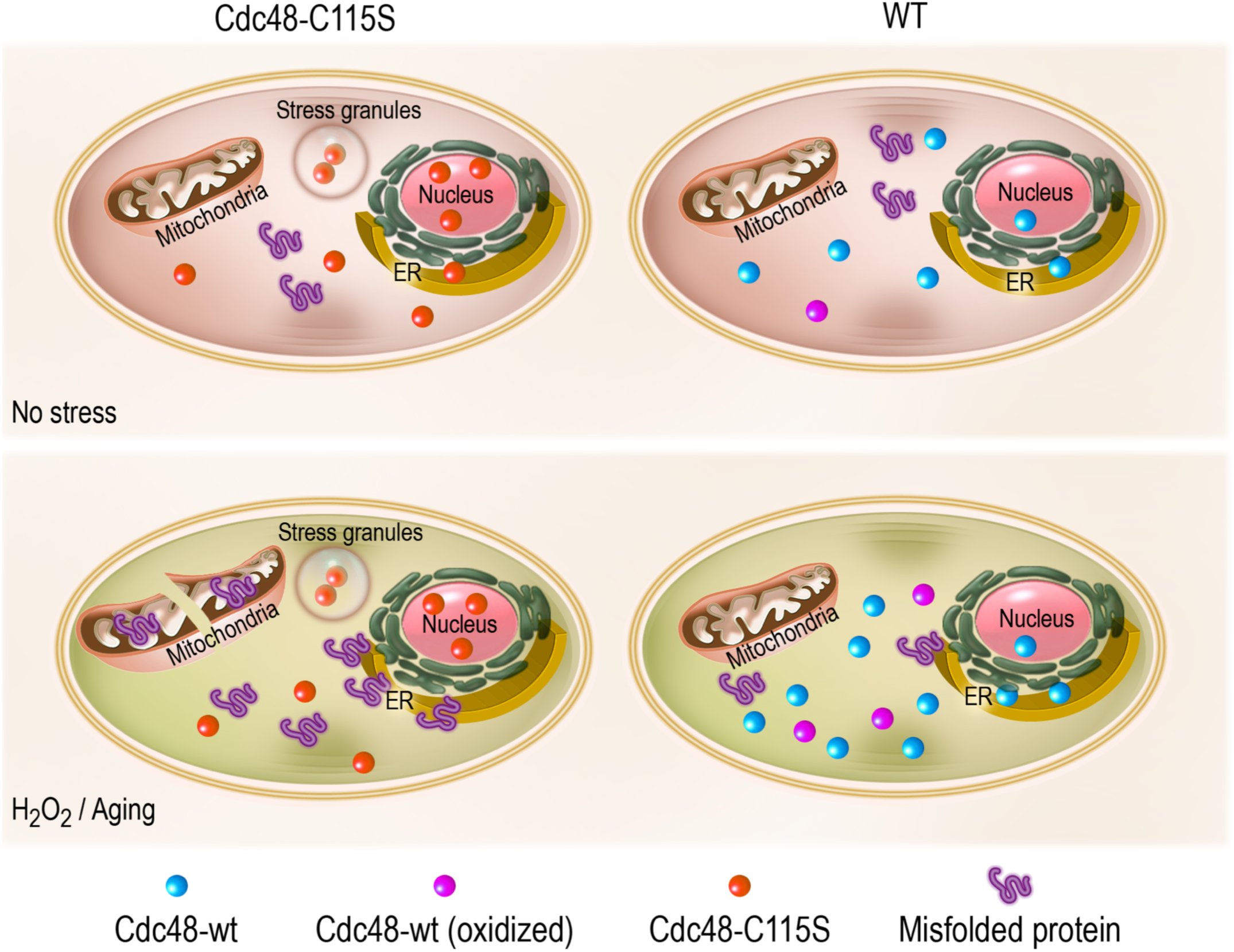
Proposed model of Cdc48 behavior under normal and stress conditions. In the wt (right), the fairly compact Cdc48 (turquoise circle) may be found localized to a variety of organelles under normal growth conditions, with relatively few unfolded proteins and some oxidized variants (magenta circle). During oxidative stress or chronological aging, the cell accumulates more unfolded proteins and thus Cdc48 is recruited to the ER and mitochondria and more cysteine residues are oxidized. In Cdc48-C115S (left), Cdc48-C115S (orange oval) is slightly more open than the wt and preferentially localized to the nucleus (as well as suspected stress granules). During oxidative stress or chronological aging, Cdc48-C115S localization does not change much relative to normal conditions, however unfolded proteins are able to accumulate in the cell and ER (triggering the UPR).

## Discussion

### Cdc48 is a link between redox and protein homeostasis

Cellular oxidants are the natural byproducts of cellular life and respiration, and may have a positive effect on the organism, as well as be involved in cell signaling and antibacterial defense^56^. On the other hand, imbalanced levels of oxidants can lead to overall protein misfolding, changes in membrane architecture, and cell death^57–59^. It was established that both accumulation of reactive oxygen species and protein misfolding are hallmarks of numerous neurological diseases and aging, thus tying these processes together. Recent studies in our and other labs have reported the discovery of a sub-group of proteins – redox switches – that use thiol oxidation to mediate diverse physiological functions, including protein quality control^26, 27, 60^, merging redox and protein homeostasis together.

In this study, we propose that one of the major players in protein quality control, Cdc48, has a crucial role in protecting protein health during oxidative stress and aging. Our results point to Cdc48 as having a pivotal responsibility through its highly conserved, redox-sensitive Cys115, which serves as a nuclear gatekeeper and regulates import of Cdc48 into the nucleus. When Cys115 is mutated to serine, a large fraction of Cdc48 accumulates in the nucleus and as such is not able to undergo necessary recruitment to the mitochondria and ER, thus precluding its ability to participate in the clearance of damaged proteins during the oxidative stress response. Interestingly, under normal conditions, such an imbalance in Cdc48’s localization does not lead to a dramatic impact on cell growth and proliferation. Remarkably, other stresses associated with protein misfolding such as heat shock and reducing stress did not have such a dramatic impact on cells expressing the Cdc48-C115S variant. Thus, the Cys115 guarding function is specifically crucial to conditions associated with ROS accumulation, such as external oxidative stress or age-dependent oxidation.

Surprisingly, despite extensive research into protein quality control mechanisms in the nucleus, little is known about the role of Cdc48 in this process. Gallagher et al., showed that Cdc48 is required for assisting the ubiquitin protein ligase, San1, in degrading insoluble proteins, expression of which led to import of Cdc48 into nucleus^38^. Yet it is not clear if such San1-dependent accumulation of Cdc48 in the nucleus affects redox homeostasis and survival during chronological aging.

We believe that our study marks a new opportunity to uncover the role of Cdc48’s localization in the ability of cells to cope with stress conditions that lead to protein misfolding, including oxidative stress. For this, we assume that the Cdc48-C115S variant might serve as a model protein to further explore the consequences rooted in accumulation of Cdc48 in the nucleus and perturbation in its distribution across the cell. We believe that these consequences are similar to ones seen in the Cdc48-C115S mutant including growth defects, a negative impact on the cells’ ability to respond to oxidative stress, changes in Cdc48’s interactome, mitochondrial fragmentation, and a decrease in canonic functions such as protein degradation in the ER and sterol biogenesis (Fig 7). Interestingly, the major effect of Cdc48 accumulation in the nucleus is limited to redox-associated functions rather than heat shock and reducing stress, suggesting multiple defense mechanisms which require active Cdc48. There is also a possibility that such redox-sensitivity in Cdc48 activity is defined by mutation of Cys115 per se or a combination of the C115S mutation and localization.

At this stage, we cannot rule out that other residues in the N-terminal domain, except the NLS domain, have a similar role. Extensive mutagenesis analysis should be done to investigate the functional role of the N-terminal domain and the NLS motif specifically in maintaining redox homeostasis.

Nevertheless, we believe that the Cdc48-C115S variant will advance our understanding of PQC mechanisms during normal and proteotoxic conditions, especially due to the fact there are very few Cdc48 inhibitors which can be used in yeast. A majority of p97/VCP inhibitors are not applicable in the yeast system, therefore an additional variant with distinct behavior is an asset in the growing field of Cdc48 research, including cell cycle progression, autophagy, protein quality control, membrane architecture, and organelle biogenesis.

### Role of the N-terminal domain in Cdc48 activity and localization

Multifunctionality of Cdc48 is encoded in its structurally flexible and highly conserved N-terminal domain, which contains the nuclear import domain^28^ and also serves as anchor domain for multiple interactions with co-factors. Therefore, it is not surprising that the overwhelming majority of VCP mutations associated with diseases are on highly conserved residues (conserved also in yeast Cdc48) and most are located in the N-terminal domain^61, 62^. Moreover, canonic mutations known to cause pathogenic disorders such as Paget’s disease of bone, frontotemporal dementia (IBMPFD) and ALS have been identified as disrupting VCP nuclear localization^63^. Unfortunately, to the best of our knowledge, only a few studies have been done to dissect redox sensitivity of cells expressing these disease-related mutations, finding, for example, that R95G and R155C mutations in VCP lead to mild sensitivity during oxidative stress^64^. Given the fact that many of the Cdc48-related pathologies, such as ALS and IBMPFD, are associated with increased oxidation and mitochondrial dysfunction, uncovering the potential impact of N-terminal domain on redox homeostasis is influential^65^.

A majority of the findings regarding the N-terminal domain of Cdc48/p97/VCP relate to interactions with co-factors and targeting of Cdc48/p97/VCP to ERAD, MAD, or other pathways. In yeast, clearance of mitochondrial proteins is mediated by interaction between the N-terminal domain and the Vms1 Cdc48-cofactor^7^, while ERAD is mediated by Shp1 interaction. Studies in mammalian cells even show that p97/VCP is involved in recognition and degradation of oxidized misfolded proteins from mitochondria^66^. In addition, dysfunction of p97/VCP ATPase activity severely impacts mitochondrial proteostasis and mitophagy^67, 68^. Our study suggests that mutation in the N-terminal domain at Cys115 results in alteration of binding of co-factors, including Shp1, and therefore a decrease in degradation of ERAD substrates and sterol biogenesis. Analysis of Cdc48’s interactome did not detect Vms1 binding, therefore we cannot be sure that morphological changes in mitochondria originate in differential binding of Vms1, though we did find a global decrease in the mitophagy-associated protein Whi2^32^ in the Cdc48-C115S strain. Despite the fact that the observed mitochondrial fragmentation is tightly coupled with the mutant strain’s redox sensitivity and the depletion of Cdc48 in cytosol, the specific mechanism is unknown as of yet.

Interestingly, our mass spectrometry analysis of Cdc48-C115S suggests that Cdc48 either interacts or influences the abundance of antioxidants within the cell (e.g., thioredoxin proteins, Trx1, Trx2, catalase, superoxide dismutase Sod1). Most probably, these are indirect interactions, which either reflect potential misfolded substrates of Cdc48 or proteins necessary for restoring the redox status of protein thiols among Cdc48-associated proteins. We may thus speculate that Cys115 is a sensor for cellular oxidation, such that its oxidation stabilizes Cdc48’s structure (as suggested by MD) and maintains antioxidant proteostasis. Additional extensive structural studies should doubtlessly be conducted to examine this phenomenon in more detail. Unfortunately, many attempts to obtain purified full-length Cdc48 variants have failed at our hands. Therefore, at this stage, we are unable to investigate the effect of the 115 position, as well as oxidation of other cysteine residues on Cdc48’s structure to the desired degree.

### Reversible modification of Cysteine 115

While there are several potential explanations for how Cys115’s modification impacts Cdc48’s activity, the link to oxidation sensitivity strongly suggests that Cys115 undergoes some manner of oxidation-based modification. Modification screens have identified Cys115’s homolog in the human p97 (Cys105) as undergoing palmitoylation^69^, which may regulate Cdc48’s association with membranes of the ER and mitochondria. Unfortunately, we were unable to identify this modification in our own mass spectrometry analyses of yeast Cdc48, using diverse methods for sample preparation and analysis. We did, however, identify Cys115 in both the oxidized and reduced forms at both 24h and 48h, while aged cells (48h) had increased fractions of oxidized cysteine 115. This supports the possibility of a reversible thiol modification at later stages of chronological aging and is in line with the redox proteomic analysis of aging yeast cells cultivated for several days at normal (2% glucose) and caloric restriction (0.5% glucose) conditions^25^. In this and our study, differential thiol-trapping was used to dissect the oxidation profile of proteins, which is restricted to detection of specifically reversible thiol modifications, such as disulfide bonds and sulfinic acid. Based on our structural model as well as homolog structures of human VCP, there is no neighboring cysteine that might be able to form an intra-disulfide bond. We cannot rule out the possibility of inter-disulfide bonds, however our attempts to detect such species using mass spectrometry did not reveal positive outcomes. Due to the large surface accessibility of the cysteine 115 residue (Fig.1, S1) we assume that sulfinic or higher oxidation modifications (e.g., sulfonic) most probably take place. Oxidation of Cys115 to sulfinic and sulfonic acids might provide an additional negative charge, which could potentially alter polar and electrostatic interactions with the NLS and Ubx-binding domains. Further quantitative methods would need to be applied and developed to identify both modification type (reversible or non-reversible) and kinetics, only after which a full picture of Cdc48’s redox-regulated working cycle can emerge.

## Acknowledgements

We are extremely grateful to the director of DNA manipulation unit at the Weizmann Institute of Science, Yoav Peleg for obtaining the CDC48-GFP constructs. We thank Einat Zalckvar for helpful consultations and providing the yeast strains and constructs. We are grateful for the financial support from the Israel Science Foundation (grant number: 1765/13 and 1537/18 for DR and 162/17 For EP), Human Frontier Science program (CDA00064/2014), the US-Israel Binational Science Foundation (grant number: 2015056), the Legacy Heritage Biomedical Science Partnership (1649/16), DIP “MitoBalance” grant (for MS). MS is an incumbent of the Dr. Gilbert Omenn and Martha Darling Professorial Chair in Molecular Genetics.

## Author contributions

Conceptualization: D.R, M. R., O. Y., E. P., T. R., and M. S.; Formal analysis: M. R., O. Y., E. S. B, and D. R.; Investigation: M. R., O. Y., Y. Y., E. S. B., R. I., and N. M-B.; Resources: N. S. and R. F.; Writing – original draft: M. R. and D. R.; Writing – review & editing: M. R., E. S. B., N. M-B., N. S., E. P., M. S., and D. R.; Supervision: D.R, I. T. A., E. P., T. R., and M. S..; Funding acquisition: D.R, E.P, M.S.

## Supplementary figures and tables

**Figure S1.**
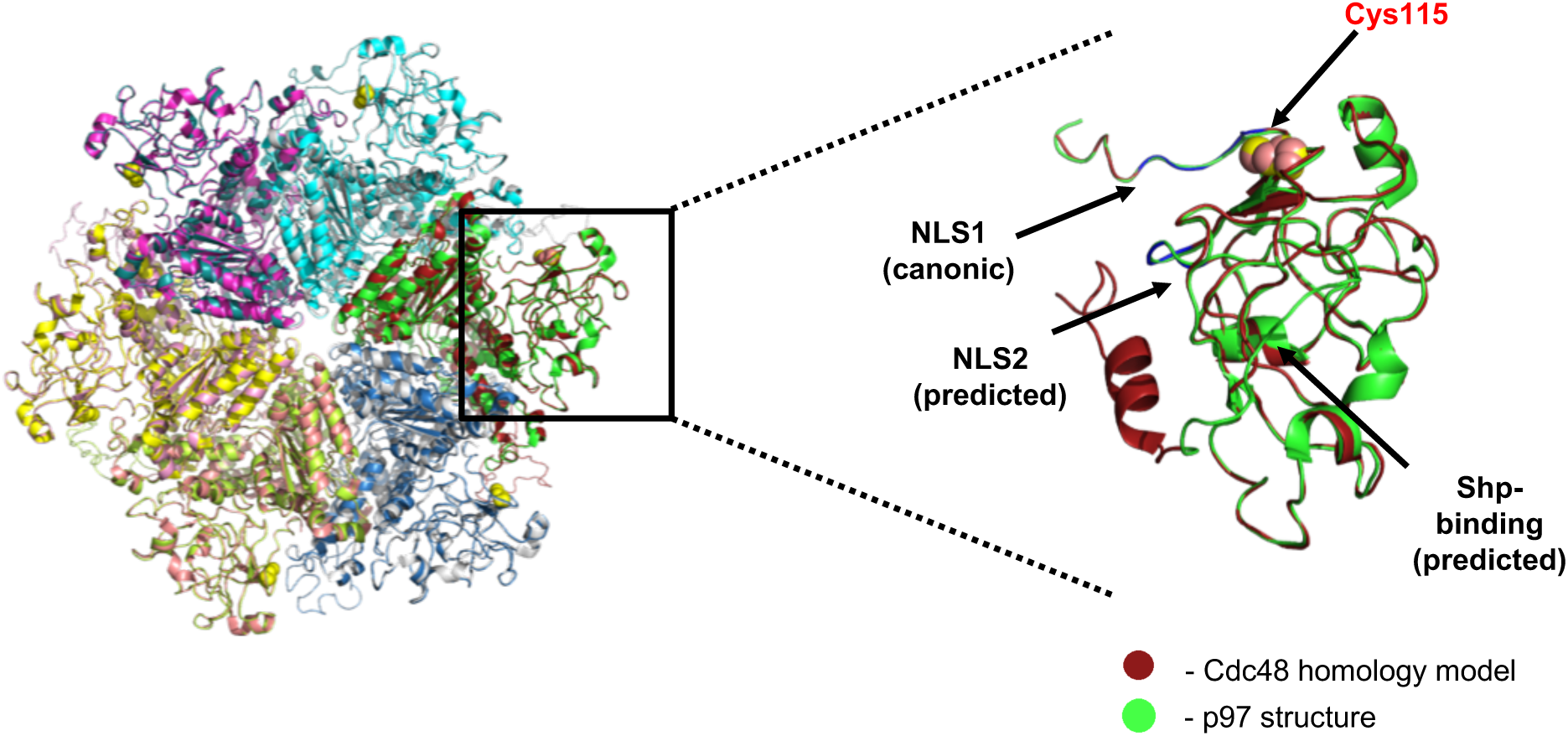
Alignment of p97 structure (PDB: 5FTN) with the homology model of Cdc48 (as shown in Fig. 1D), zoomed in on the alignment of the N-domain.

**Figure S2.**
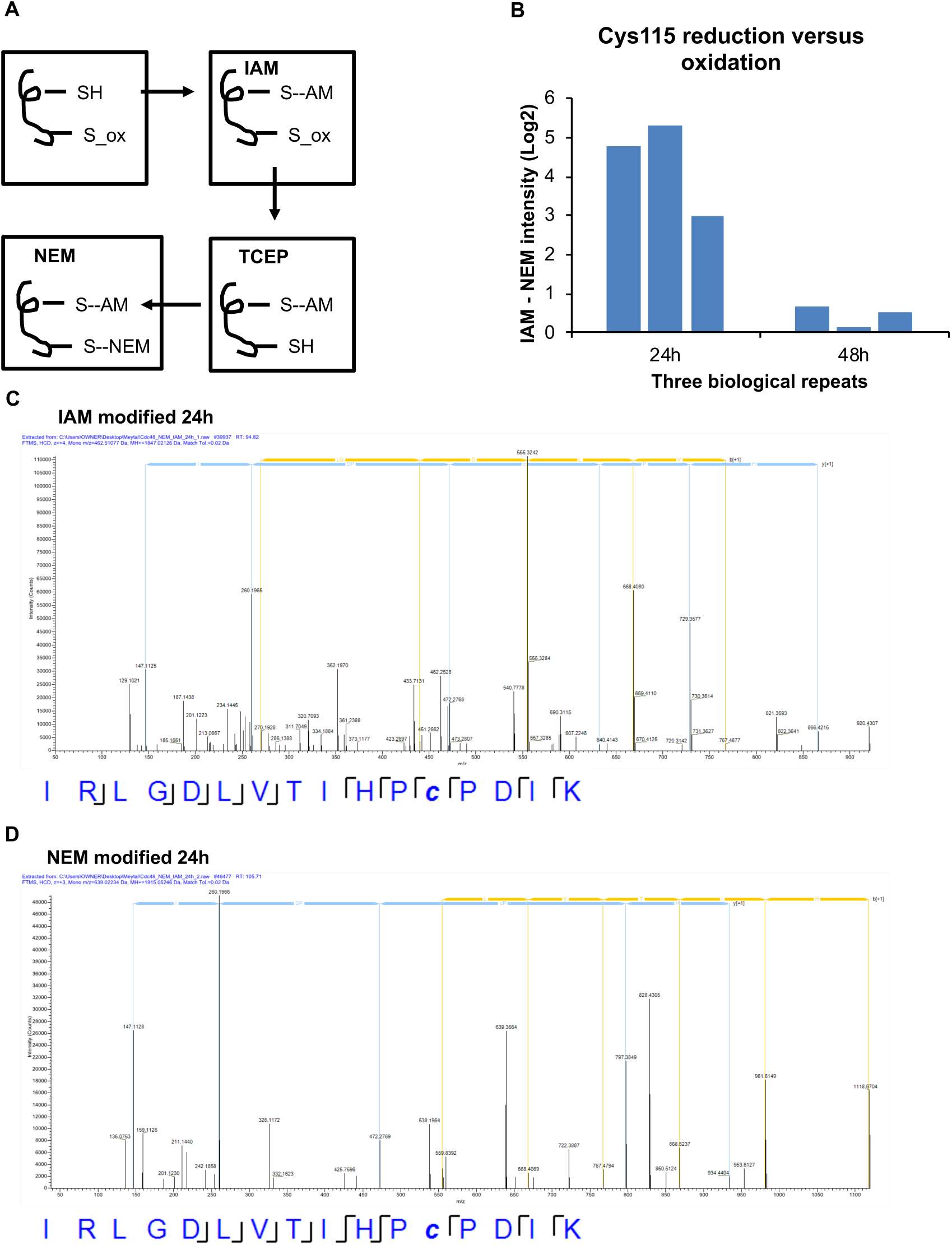
(A) NEM-IAM workflow. Proteins are initially treated with iodoacetamide (IAM), which binds any reduced cysteine residue while oxidized cysteines remain unchanged. Samples are then reduced using tris(2-carboxyethyl)phosphine (TCEP) to reduce remaining cysteine residues, after which they are treated with N-ethylmaleimide (NEM), labeling the formerly oxidized cysteine residues. (B) Difference in log2 intensity between IAM- and NEM-labeled peptides in the mass spectrometer. Cys115 has a higher intensity of IAM-labeling at 24h as compared with 48h, in three biological replicates (various peptide repeats). (C) Representative spectrum of IAM-modified peptide containing Cys115 at 24h. (D) Representative spectrum of NEM-modified peptide containing Cys115 at 24h.

**Figure S3.**
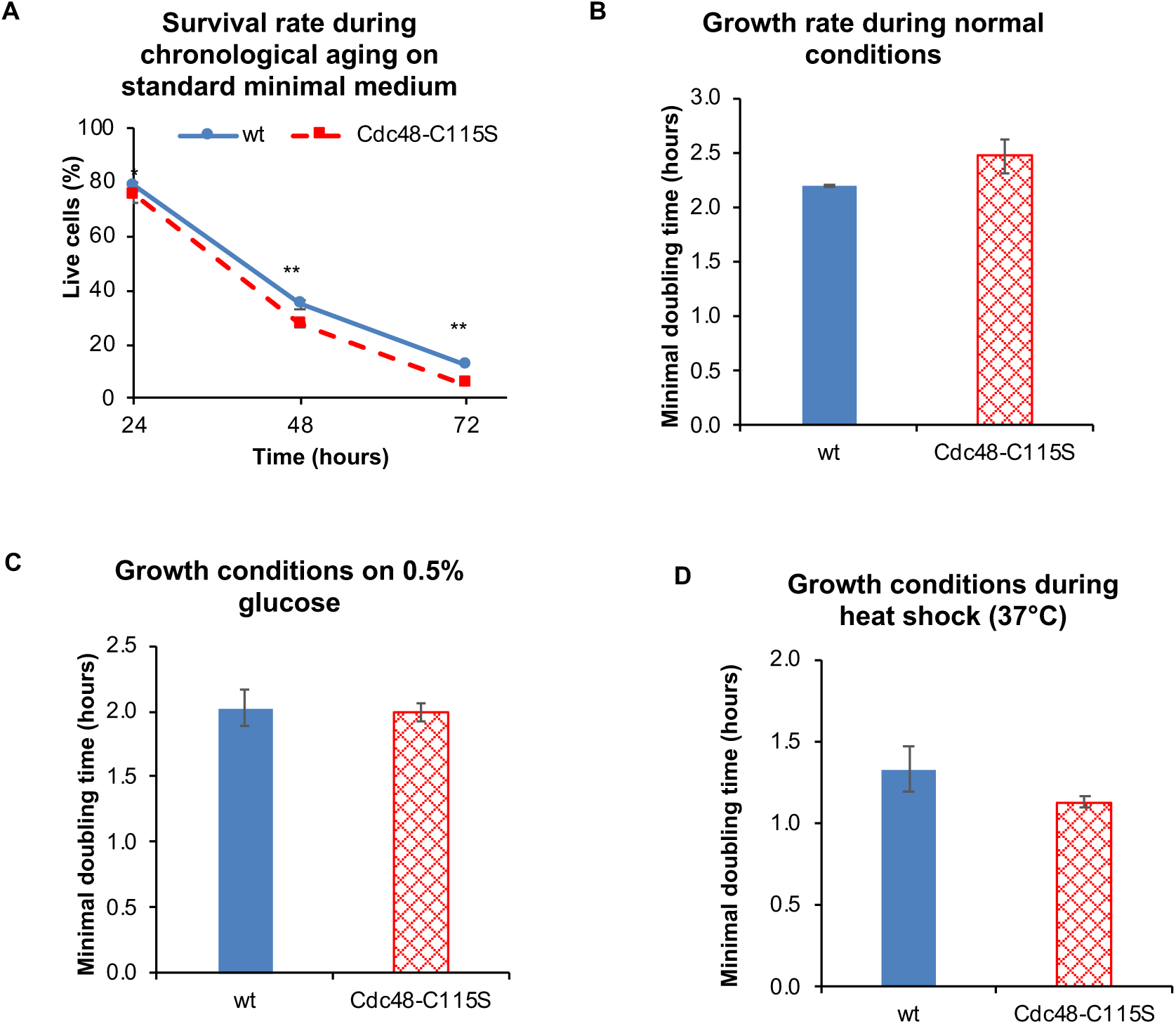
(A) Survival of living cells expressing either the wt or Cdc48-C115S variant, grown on standard minimal medium, with Cdc48-C115S again displaying lower vitality. (B) Cdc48-C115S has slightly impaired growth under normal conditions as compared to the wt. (C) Growth on 0.5% glucose is not significantly different between wt and Cdc48-C115S. (D) Growth during heat shock (37°C) is not significantly different between wt and Cdc48-C115S.

**Figure S4.**
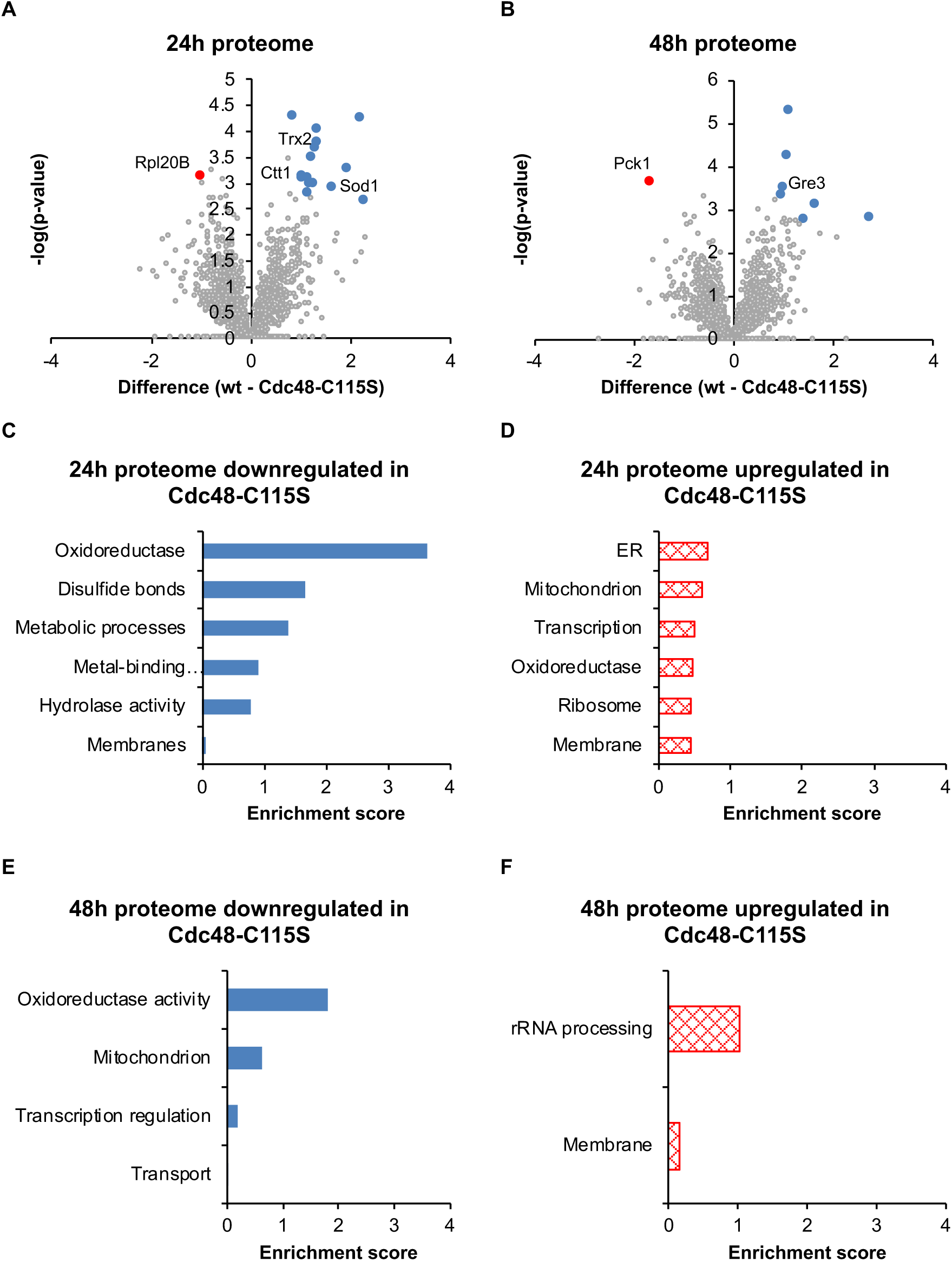
Mutation of Cdc48-C115S does not severely alter the cellular proteome. (A-B) Volcano plots of differentially regulated proteins between the wt and Cdc48-C115S at 24h and 48h (respectively), according to an FDR of 0.05. Proteins relatively downregulated in Cdc48-C115S are labeled blue, upregulated are red. Proteins that were not expressed at all in one group are unlabeled, but analyzed. (C-F) Functional enrichment analysis of all differentially regulated proteins, according to regulation and chronological age. The only major functional group to significantly change is a downregulation of oxidoreductase function in Cdc48-C115S at both 24h and 48h.

**Figure S5.**
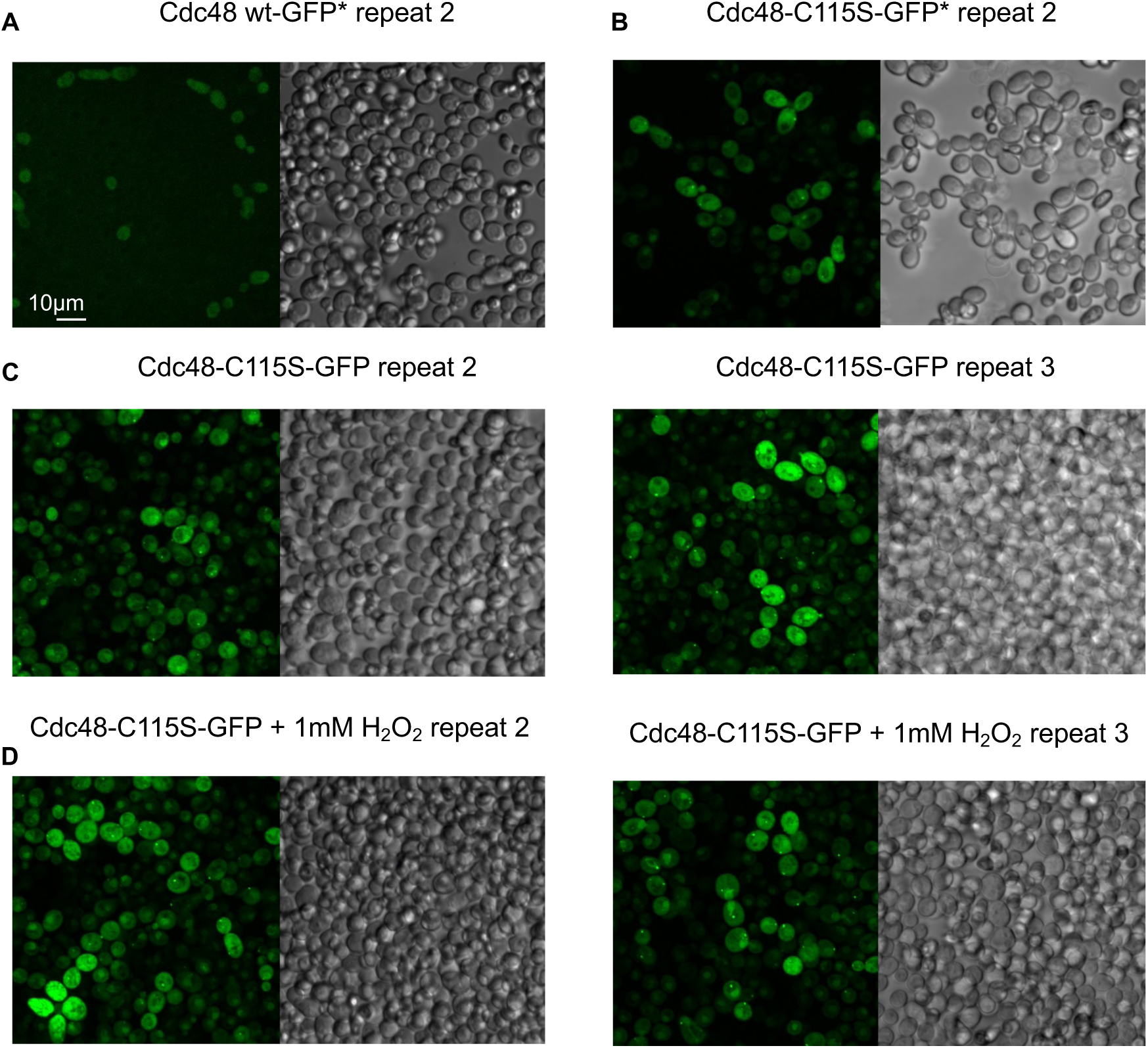
Corresponding repeat fields for Fig. 4. (A) Biological repeat field of wt-GFP*. (B) Biological repeat field of Cdc48-C115S-GFP*. (C) Biological repeat fields of Cdc48-C115S-GFP. Repeat 3 is from a separate experiment under the same growth and imaging conditions. (D) Biological repeat fields of Cdc48-C115S-GFP treated with 1mM H_2_O_2_. Repeat 3 is from a separate experiment under the same growth and imaging conditions.

**Figure S6.**
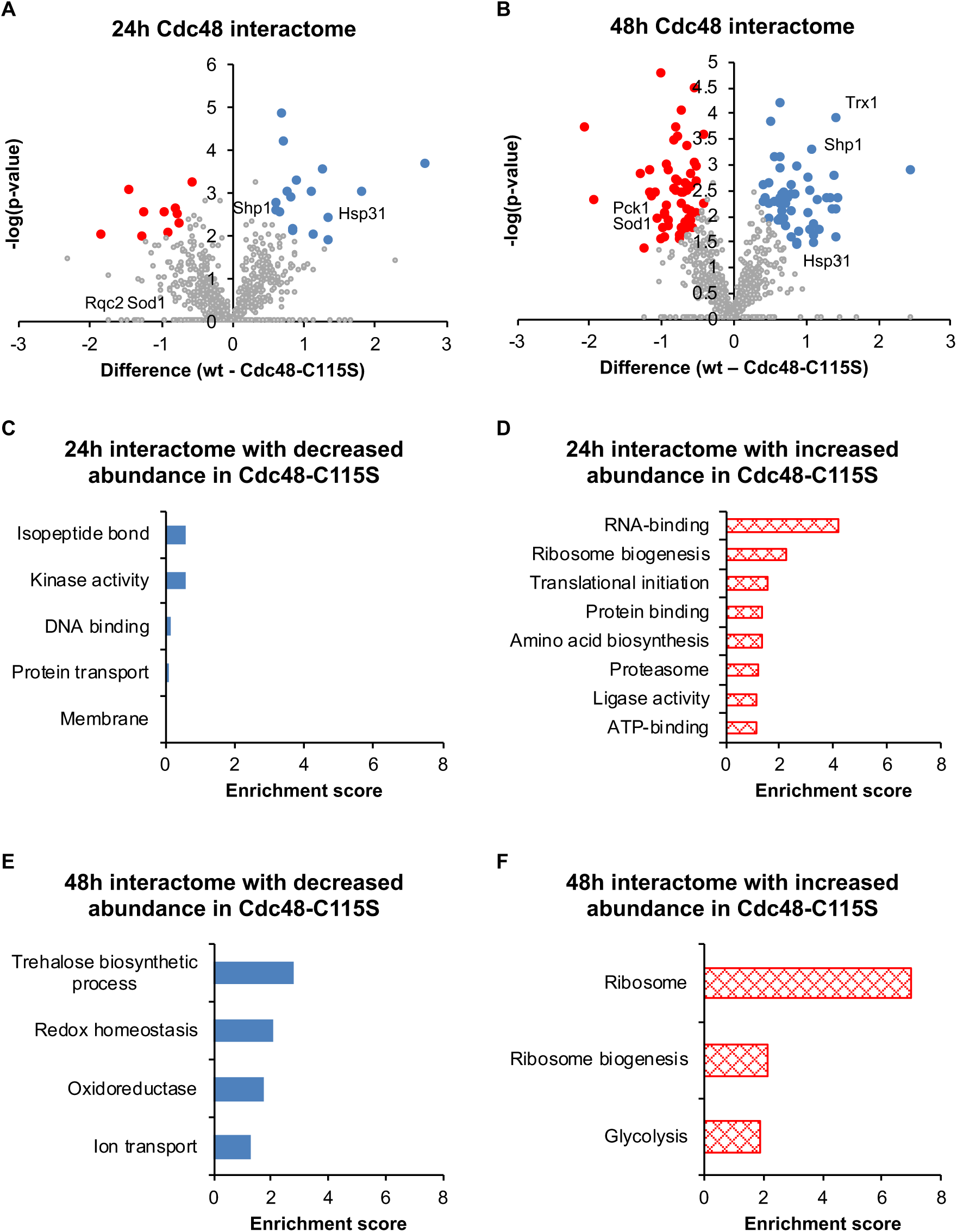
Mutation of Cdc48-C115S significantly alters Cdc48’s interactome. (A-B) Volcano plots of differentially regulated proteins between wt and Cdc48-C115S’s interactome at 24h and 48h (respectively), according to an FDR of 0.05. Proteins with relatively decreased abundance in Cdc48-C115S are labeled blue, increased abundance are red. Proteins that were not expressed at all in one group are unlabeled, but analyzed. (C-F) Functional enrichment analysis of all differentially regulated proteins, according to regulation and chronological age. Ribosomal and RNA-associated functions were strongly enriched in the Cdc48-C115S’s interactome at both 24h and 48h, while redox-associated functions were decreased at 48h.

**Figure S7.**
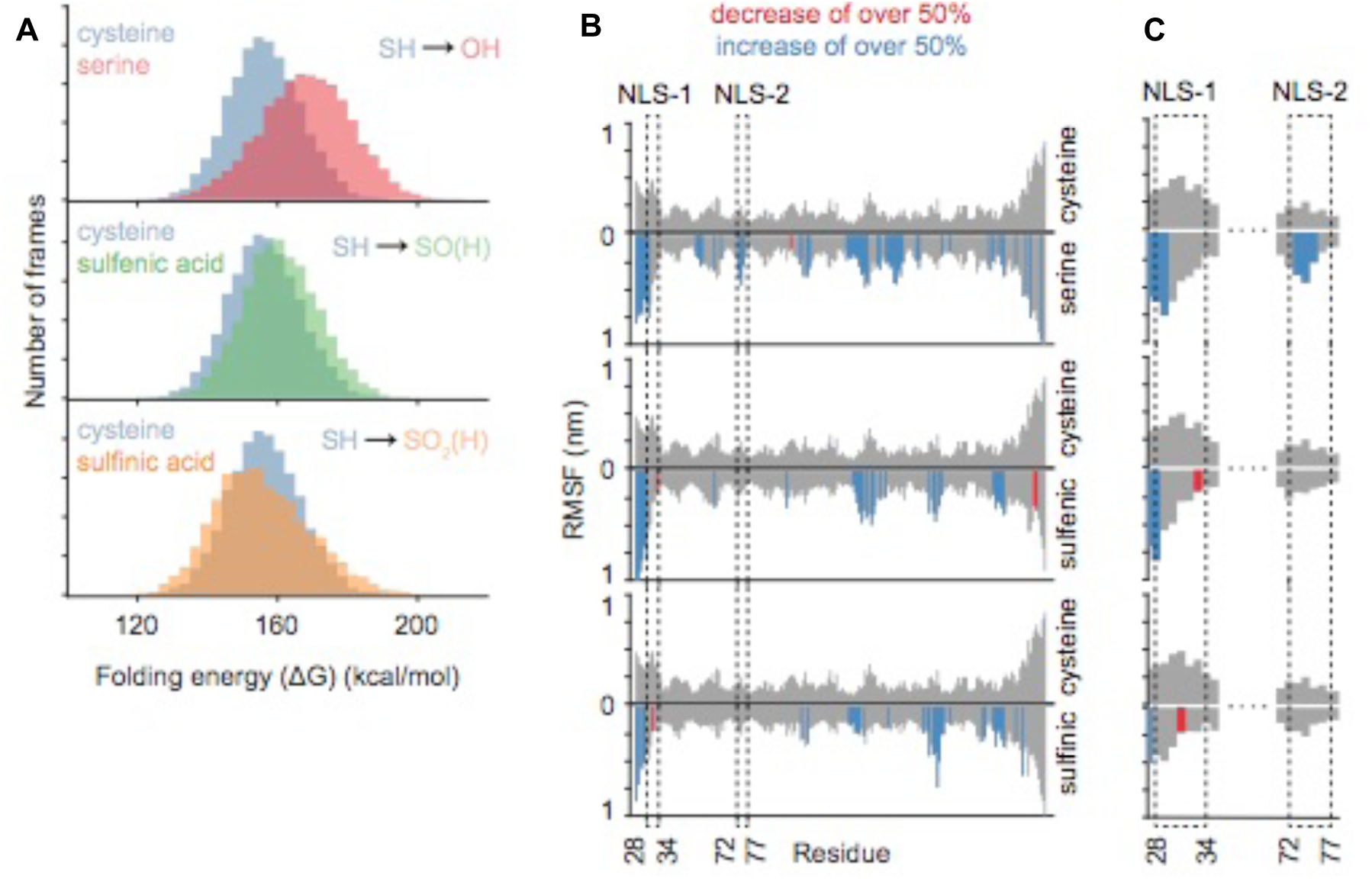
(A) Histogram of Cdc48 folding energy of the wt model (blue) compared to the three mutant models (serine mutant in red and oxidized cysteine side chains in green and yellow). (B) Root mean square fluctuation (RMSF), or standard deviation, of each residue over the course of the production simulation from 10ns to 100ns. Blue or red bars indicate positions that have an increase or decrease, respectively, in fluctuations by over 50% compared to wt. (C) A zoom in on the NLS-1 and NLS-2 regions of the RMSF graph.

**Figure S8.**
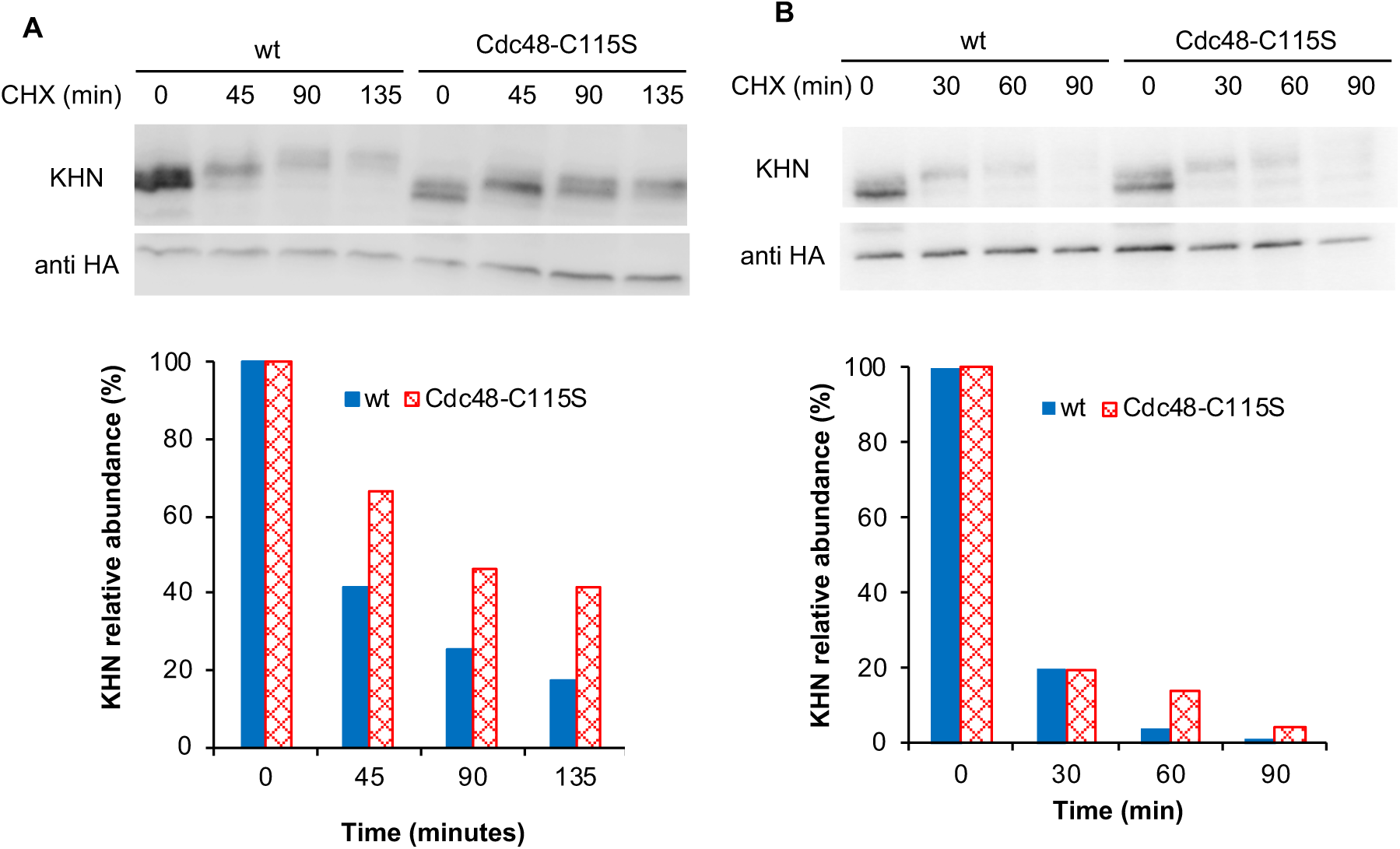
Repeat experiments of KHN degradation using pulse-chase, corresponding with Fig. 6C-D, with corresponding quantification.

**Figure S9.**
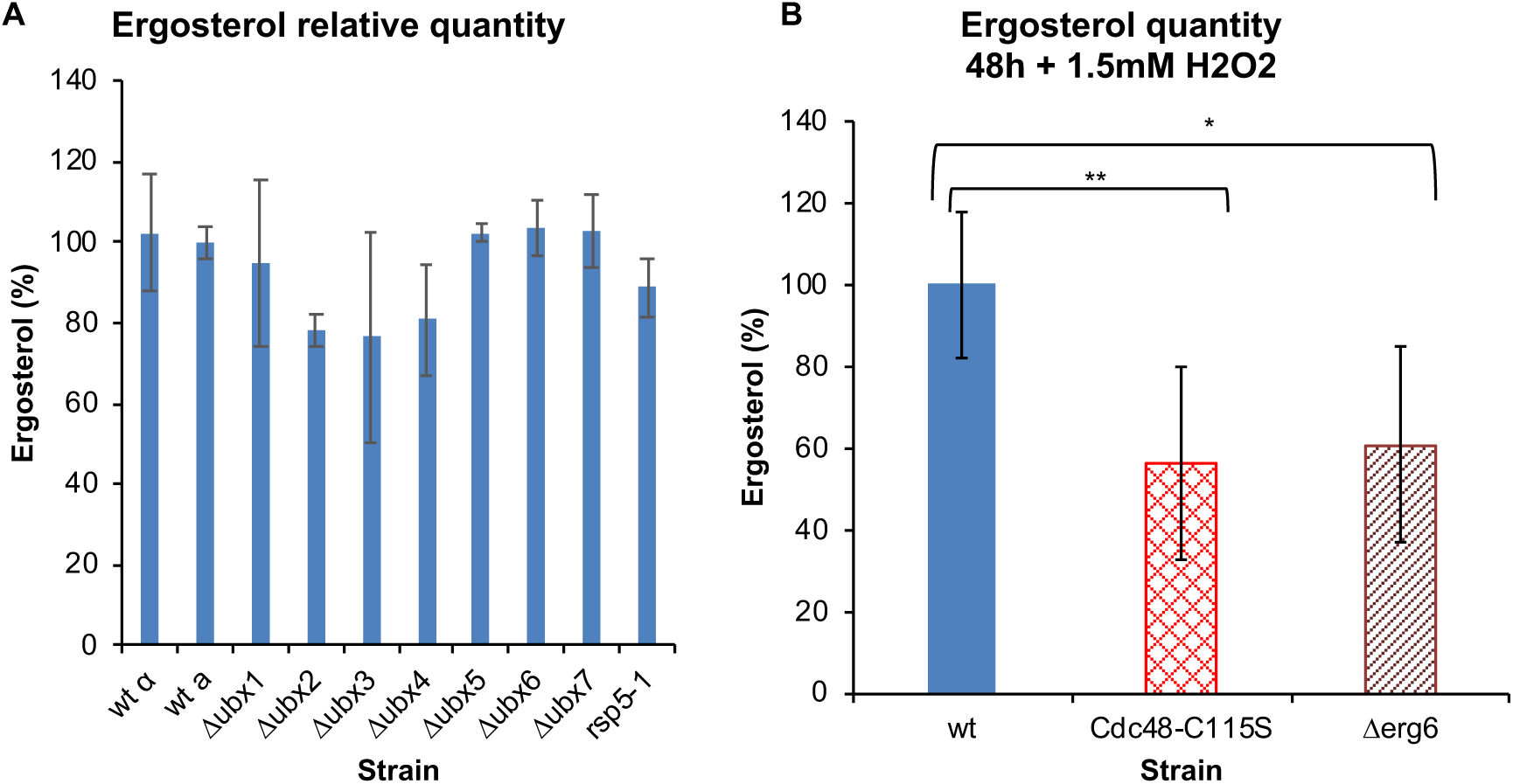
Ergosterol quantity at 48h during oxidative stress. Cdc48-C115S has a significantly reduced quantity of ergosterol compared to the wt, comparable to the ergosterol-defective Δerg6 strain.

## Tables

**Table S1.** Cdc48 homology sequence identity in yeast (*Saccharomyces cerevisiae*), Arabidopsis (*Arabidopsis thaliana*), drosophila (*Drosophila melanogaster*), human (*Homo sapiens*), and zebrafish (*Danio rerio*) homologs (% identity).

| % identity | *S. cerevisiae* | *A. thaliana* | *D. melanogaster* | *H. sapiens* | *D. rerio* |
| --- | --- | --- | --- | --- | --- |
| *S. cerevisiae* | 100 | 67 | 69 | 69 | 69 |
| *A. thaliana* | 67 | 100 | 75 | 77 | 77 |
| *D. melanogaster* | 69 | 75 | 100 | 83 | 83 |
| *H. sapiens* | 69 | 77 | 83 | 100 | 97 |
| *D. rerio* | 69 | 77 | 83 | 97 | 100 |

Tables S2-S11 are attached separately

**Table S2.** All proteins in the wt and Cdc48-C115S all-cell proteomes during 24h.

**Table S3.** All proteins in the wt and Cdc48-C115S all-cell proteomes during 48h.

**Table S4.** Significantly differentially regulated proteins between the wt and Cdc48-C115S all-cell proteomes during 24h.

**Table S5.** Significantly differentially regulated proteins between the wt and Cdc48-C115S all-cell proteomes during 48h.

**Table S6.** All proteins in the wt and Cdc48-C115S interactomes following co-immunoprecipitation during 24h.

**Table S7.** All proteins in the wt and Cdc48-C115S interactomes following co-immunoprecipitation during 48h.

**Table S8.** Significantly differentially regulated proteins between the wt and Cdc48-C115S interactomes during 24h, following removal of background interactions.

**Table S9.** Significantly differentially regulated proteins between the wt and Cdc48-C115S interactomes during 48h, following removal of background interactions.

**Table S10.** Significantly differentially regulated proteins between the wt and Cdc48-C115S interactomes during 24h, without removal of background interactions.

**Table S11.** Significantly differentially regulated proteins between the wt and Cdc48-C115S interactomes during 48h, without removal of background interactions.

## Materials and Methods

### Cdc48 strain preparation

Plasmids containing both Cdc48-wt and Cdc48-C115S were prepared C-terminally tagged with either FLAG and HIS-tag sequences. Plasmids are based on pRS415(LEU2), under the TEV promoter. These plasmids were then transformed according to protocol modified from^1^ alongside deletion of the chromosomal Cdc48 sequence through a diploid strain containing *Δcdc48*, with viable clones selected following sporulation. Clones were confirmed by sequencing. GFP-tagged plasmids are based on pRS416(URA3). These plasmids were transformed into existing Cdc48-FLAG strains *(Δcdc48* with plasmid-based *CDC48*). Certain strains lost their ability to grow on media lacking Leu, indicating existence of the URA plasmid only. Strains with both resistances are labeled GFP*.

### Bioinformatics, homology modeling, and structural analysis

Multiple sequence alignment was conducted using ClustalOmega^2^, with sequences acquired from the Uniprot database^3^. Structural data was based on solved protein structures 5C1B^4^ and 5FTN^5^ acquired from the Protein Data Bank, with all images done in PyMOL (multiple versions), Schrödinger, LLC. These PDB entries were used as templates for Cdc48 homology modeling under SWISS-MODEL^6^. The model based on 5C1B was created following a search for compatible Cdc48-sequence candidate templates, resulting in a single monomer. The homo-hexamer was prepared by separate alignment of p97-5C1B with each chain redefined to match the corresponding monomer on p97. The model based on 5FTN was created by uploading 5FTN’s structure as a target template. All settings were set to default values. Image in Fig. 1B used according to Creative Common License: Somersault1824.

### Growth mediums

We used three different growth mediums at different points throughout this research. For general, non-selective growth, we used a rich growth medium YPD: 10g yeast extract, 20g peptone (2%), 20g glucose (2%), 4ml 1% adenine, dissolved in 1 liter DDW, autoclaved. Selective growth on a partially synthetic, casein-supplemented medium (semi-rich) was on a casamino acid rich medium: 0.017% yeast nitrogen base (w/o amino acids and ammonium sulfate), 0.5% ammonium sulfate, 2% glucose (unless otherwise specified), 0.2% casamino acid mix, 0.000004% Trp, 0.000005% Thr (amino acids to excess; subject to minor variation), 4ml 1% adenine in 1 liter DDW. The medium may also be supplemented with 0.5% uracil, depending on strain selection. For minimal selective medium, we used a synthetic medium: 1.7g yeast nitrogen base (without amino acids or ammonium sulfate), 5g ammonium sulfate (0.5%), 20g glucose (2%), 10ml selective amino acid mix x100 (all amino acids somewhat to excess), 4ml 1% adenine, 4ml 1% uracil, in 1 liter DDW, autoclaved. Unless otherwise noted, all experiments were conducted on the partially synthetic, casein-based medium.

### Co-immunoprecipitation

Samples were grown on 2-4ml synthetic medium supplemented with casein amino acid, then diluted to 0.2 O.D._600_ on 5-10ml medium and grown for varying degrees of time. Cells were harvested and broken in “Extraction buffer” (0.5ml Tris HCl pH 7.5 buffer with protease inhibitors) (all stages conducted on ice or at 4°C), using glass beads and 4 cycles of disruption (20s pulse, 30s on ice). Cells were then diluted with 260µl Extraction buffer and 40µl 20% Triton in order to solubilize membranes and centrifuged at low speed (0.4 x g) for 5 minutes in order to clear broken cells. Supernatant was then centrifuged at 13,300 x g for 30 minutes, after which the supernatant was transferred to a new 2ml tube. Supernatant was diluted 1:1 with “Resuspension buffer” (50 mM Tris-HCl, pH 7.5, 200 mM NaAc, 10% glycerol) and protease inhibitors. Magnetic beads were prepared in buffer and added to each sample and incubated overnight at 4°C under constant agitation. Next morning, samples were cleared of the supernatant and beads washed several times before proceeding to sample preparation for mass spectrometry (as described previously).

### Monitoring yeast growth using plate reader

Growth curves were conducted on diluted yeast samples (0.1-0.2 O.D._600_) using a TECAN Infinite 200 PRO series. Treatment with reagents (i.e. H_2_O_2_, DTT) was conducted within the wells, with the reagent added to the growth medium into which the yeast cultures were diluted. Yeast incubation and OD_600_ measurements, as well as minimal doubling times, were analyzed using MDTCalc^7^. Minimum doubling time was assessed by first calculating the log10 value of each point during logarithmic growth, and then measuring the slope between each set of consecutive points. The value of this slope was then inverted (representing the minimal time it took the sample point to “double”), providing the minimal doubling time value.

### Microscopy

Yeast were images using the FV-1200 confocal microscope (Olympus., Japan), using a 60X/1.42 oil objective. GFP was excited with 488nm laser and emission was collected 500-540nm, a DIC image was also taken. For mitochondrial labeling, cells were grown to mid-log phase (after dilution). Treatment with 1mM H_2_O_2_ was for 1 hour. For the Cdc48-GFP imaging, cells were grown to 24h growth (after dilution). Treatment with 1mM H_2_O_2_ was for 1 hour. Cells were counted by hand. Figures were prepared using ImageJ. Fluorescent images were compiled through Z project per maximum intensity, while DIC images through Z project per average intensity.

### Mass spectrometry and proteomic analyses

For all-proteome analysis, 5mL of stationary phase yeast cultures (24h, 48h) were lysed using 300µl 0.2M NaOH and resuspended using a lysis buffer (100mM DTT [15.425mg DTT], 100µl 1M Tris HCl pH 7.5, 100µl SDS 20%, complete with DDW to 1ml). Samples were then diluted using 400µl of a urea buffer (8M urea in 0.1M Tris HCl pH 8.5), loaded onto a filter, and centrifuged for 10 minutes at 12,000g following the standard FASP protocol^8^. This process was repeated three times, with flow-through discarded, after which samples were incubated in the dark for 60 minutes with 0.5M iodoacetamide and urea buffer (final iodoacetamide concentration of 0.05M), under constant agitation (350rpm, 25°C). Samples were then washed again three times with urea buffer and twice with digestion buffer (10% ACN, 25mM Tris HCl pH 8.5), then centrifuged at 12,000g for 8 minutes. Filters were transferred to a new collection tube, suspended in 300µl of digestion buffer with 1µl of trypsin (Promega), mixed for 1 minute at 600rpm, and left overnight at 350rmp, 37°C. Following digestion, samples were centrifuged for 10 minutes at 12,000g. Cdc48 interactome samples were similarly prepared, following lysis and interactome pull-down according to the Co-IP method described previously.

The peptide concentration was determined, after which the peptides were loaded onto stage tips in equal amounts. Stage tips were activated using 100µl MS-grade methanol (100% MeOH) and centrifuged for 2 minutes at 2,000g, after which they were cleaned with 100µl of elution buffer (80% ACN, 0.1% formic acid) and centrifuged again for 2 minutes at 2,000g. The stage tips were returned to their hydrophilic state by suspension in 100µl of buffer A (0.1% HPLC-grade TFA) and centrifugation at 2,000g for 2 minutes, repeated once. 10-30µg of protein was then loaded per stage tip (as per protein preparation above), and centrifuged at 1,000g for 2 minutes. Proteins were then washed twice with 100µl buffer A at 1,000g for 2 minutes and transferred to a new collection tube. Peptides were eluted using 60µl buffer B (80% ACN, 0.1% HPLC-grade TFA) centrifuged at 250g for 2 minutes, and another 30µl buffer B centrifuged at 250g for 2 minutes. Samples were then dried using a SpeedVac for 24 minutes at 1,300rpm at 35°C, after which they were dissolved in 6-12µl of buffer A and prepared for tandem mass spectrometry analysis.

### Nano-LC-MS/MS analysis

The peptides were injected into a Nano Trap Column, 100 µm i.d.×2 cm, packed with Acclaim PepMap100 C18, 5 µm, 100Å (Thermo Scientific) for 8 min at flow 5ul/min, and then separated on a C18 reverse-phase column coupled to the Nano electrospray, EASY-spray (PepMap, 75mm x 50cm, Thermo Scientific) at flow 300 nl/min using an Dionex Nano-HPLC system (Thermo Scientific) coupled online to Orbitrap Mass spectrometer, Q Exactive Plus (Thermo Scientific). To separate the peptides, the column was applied with a linear gradient with a flow rate of 300 nl/min at 45°C: from 1 to 35% in 100 min, from 35 to 55% in 43 min, from 55 to 90% in 5 min, and held at 90% for an additional 30 min, and then equilibrated at 1% for 20 min (solvent A is 0.1% formic acid, and solvent B is 80% acetonitrile, 0.1% formic acid). The Q Exactive was operated in a data-dependent mode. The survey scan range was set to 200 to 2000 m/z, with a resolution of 70,000 at m/z. Up to the 12 most abundant isotope patterns with a charge of ≥2 and less than 7 were subjected to higher-energy collisional dissociation with a normalized collision energy of 28, an isolation window of 1.5 m/z, and a resolution of 17,500 at m/z. To limit repeated sequencing, dynamic exclusion of sequenced peptides was set to 60 s. Thresholds for ion injection time and ion target value were set to 70 ms and 3×10^6^ for the survey scans and to 70 ms and 10^5^ for the MS/MS scans. Only ions with “peptide preferable” profile were analyzed for MS/MS. Data was acquired using Xcalibur software (Thermo Scientific). Column wash with 80% ACN for 40 min was carried out between each sample run to avoid potential carryover of the peptides.

### Data Analysis and statistics of the proteomic data

For protein identification and quantification, we used MaxQuant software^9^, versions 1.5.3.30 and 1.6.3.3. We used Andromeda search incorporated into MaxQuant to search for MS/MS spectra against the UniProtKB database of Saccharomyces cerevisiae proteome, (Uniprot release, Aug 2016). The identification allowed two missed cleavages. Enzyme specificity was set to trypsin, allowing N-terminal to proline cleavage and up to two miscleavages. Peptides had to have a minimum length of seven amino acids to be considered for identification. Carbamidomethylation was set as a fixed modification, and methionine oxidation was set as a variable modification. A false discovery rate (FDR) of 0.05 was applied at the peptide and protein levels. An initial precursor mass deviation of up to 4.5 ppm and fragment mass deviation up to 20 ppm were allowed. Only proteins identified by more than 2 peptides were considered. To quantify changes in protein expression we used the label-free quantification (LFQ) using the MaxQuant default parameters. For statistical and bioinformatic analysis, we used Perseus software (http://141.61.102.17/perseus_doku/doku.php?id=start). For functional enrichment analysis, the DAVID webserver^10^ was used, as well the UniProt database and the Yeast Genome Database^11^ for both localization and additional functional enrichment analysis. Protein localization was determined using the Loqate database^12^, supplemented with data from the Yeast GFP database^13^.

### NEM-IAM preparation and data analysis

Samples were prepared using co-immunoprecipitation method as previously described, followed by modified FASP protocol. All buffers for both co-IP and FASP were first placed in an anaerobic chamber to reduce accidental oxidation during sample preparation. Samples were treated with 0.05M IAM as in the previously described FASP protocol, after which they were washed twice with urea and treated with 0.005M TCEP. Samples were washed twice more with urea and then treated with 0.05M NEM. All treatments were carried out for 60 minutes under 350rpm shaking at 25°C. Remaining FASP, stage tips, and mass spectrometry protocols followed as previously described. Peptide identification, quantification, and spectrum visualization was done using Proteome Discoverer (Thermo Fisher Scientific), identifying all peptides containing the cysteine residue of interest. The intensity of each identified peptide with either the IAM or NEM modification (high confidence) was summed. The difference in the log2 values of the summed intensity was represented graphically per individual biological repeat.

### Unfolded protein response assay

Strains were transformed with a plasmid provided by Maya Schuldiner’s lab, expressing a GFP-labeled unfolded protein response element. The transformation was conducted according to homologous recombination, according to the protocol from^1^. Cells were washed with 0.1M LiAc with sorbitol, then incubated at 42°C for 40 minutes with 100µl 0.1M LiAc, 20µl freshly boiled salmon sperm DNA mix (5mg/ml), 20µl PCR product (DNA), and 700µl fresh 40% PEG 3350 with LiAc and 10 x TE, resuspended in sterile water, plated, and left to recover for 3-4 days. GFP-expression was measured according to fluorescence using flow cytometry, with values then normalized in comparison with daily expression of the wt strain due to changes in plasmid expression during growth.

### Pulse chase assay

Yeast cells expressing a KHN-HA plasmid (provided by the Ravid lab) were grown overnight, diluted to 0.2-0.3 O.D. and grown on selective medium enriched with casein for 3-4 hours until the logarithmic phase (0.6 O.D.). Cells were then harvested and resuspended again at ∼10^7^ cells/1mL. Samples were prepared for each time point (e.g. 0, 45, 90, 135 minutes), with 100µl sodium azide 0.1M (on ice). 250µl of 10mg/ml cycloheximide (or a ratio of 1:20) was added to the cells, vortexed briefly, and incubated for relevant lengths of time before cells were lysed and prepared for Western Blot.

### Ergosterol assay

Ergosterol assays were performed per standard protocol^14^, with minor changes. Strains were grown overnight on synthetic medium as previously described and 20 OD_600_ were harvested in a 2ml flat bottom microcentrifuge tube and washed once with sterile DDW. The pellet was then suspended in 600µl of 30% alcoholic potassium hydroxide solution (3g of KOH and 3.5ml sterile DDW, brought to 10ml with 100% ethanol) and incubated in 90°C heat block under constant shaking for 2 hours. Next, 600µl of n-Heptane and 55µl DDW were added to the mix, which was subsequently vigorously mixed for 3 minutes at maximum speed in a vortex. Finally, 200µl of the upper phase was used for analysis in a 96-well plate reader (Biotek Synergy HT). Results are presented as mean ± standard deviation for data averaged from at least three independent biological repeats. To determine statistical significance, Student’s t-test or one-way ANOVA were used. Post-hoc analysis was performed with Tukey multi-comparison test. For all experiments, significance was determined by p<0.05.

### Western Blot analysis

Following lysis using 0.1M NaOH, samples were resuspended in sample buffer (Tris HCl 125mM pH 6.8, 4% SDS, 20% glycerol, 0.004% bromophenol blue), boiled at 96°C for 5-10 minutes, and centrifuged at 4,000g for 1 minute. Samples were then loaded onto an SDS-PAGE gel (10% acrylamide) and transferred to a nitrocellulose membrane using a Trans-blot Turbo System (BioRad), using a 30 minute program of 25V, 2.5A. The membrane was then treated with a blocking solution for one hour (10% skim milk in TBS with Tween 20) and incubated with the primary antibody at a ratio of 1:5,000 for 1 hour at room temperature (or overnight at 4°C) under constant motion. The membrane was then washed three times with TBST and incubated with the secondary antibody at a ratio of 1:10,000 for 30-60 minutes at room temperature under constant motion. The membrane was then briefly washed again three times, incubated for ∼1 minute with EZ-ECL reagents (Biological Industries) under constant motion at room temperature, and photographed using a Lumitron photography system. Protein band intensity was assessed using a pixel-counting program, normalized against the nonspecific HA band.

### Grx1-roGFP2 probes

The strains were transformed with Grx1-roGFP2 (kindly provided by Bruce Morgan). Transformations were carried out using a standard “One Step” protocol, modified from Chen, et al.^15^, as described in Chapter 1. Cultures were diluted to an OD_600_ of approximately 0.25 on 4-10ml to ensure fresh growth and brought to their logarithmic phase of OD_600_ of approximately 0.75, which marked “time zero” (or “log”). Samples were grown at 30°C under constant agitation. Transformations were refreshed every 2-3 weeks in an adapted version of the protocol from^16^.

### Molecular dynamics

The initial structure for the Cdc48-C115S mutant was built by replacing the sidechain sulfur with an oxygen. The initial oxidized cysteine structures were built using the Vienna-PTM webserver^17^. The simulations were performed using GROMACS 2018 with the GROMOS 54A7 force field. All four initial models were solvated in an extended simple point charge water model. Ions were added to neutralize the system charge. Initial energy minimization of these four solvated systems was performed using the steepest-descent algorithm with convergence occurring when the maximum force was less than 1000 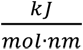. By applying positional restrains of 1000 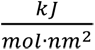 on the protein heavy atoms, followed by a two-step equilibration process with a constant number of particles, volume, and temperature (NVT), and then with a constant number of particles, pressure, and temperature (NPT), the water and ions were allowed to equilibrate around the protein. The reference temperature and pressure for equilibration were 300K and 1bar. A 100ns production MD simulation was then performed with a time step of 2 fs. The trajectory included 10,000 frames. Energy and fluctuation calculations were performed from frames representing 10ns-100ns of the production simulation. The folding energy was calculated for every relevant frame in 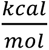 using FoldX Suite 4.0^18^.

